# Dynorphin modulates motivation through a pallido-amygdala cholinergic circuit

**DOI:** 10.1101/2024.07.31.605785

**Authors:** Qingtao Sun, Mingzhe Liu, Wuqiang Guan, Xiong Xiao, Chunyang Dong, Michael R. Bruchas, Larry S. Zweifel, Yulong Li, Lin Tian, Bo Li

## Abstract

The endogenous opioid peptide dynorphin and its receptor κ-opioid receptor (KOR) have been implicated in divergent behaviors, but the underlying mechanisms remain elusive. Here we show that dynorphin released from nucleus accumbens dynorphinergic neurons exerts powerful modulation over a ventral pallidum (VP) disinhibitory circuit, thereby controlling cholinergic transmission to the amygdala and motivational drive in mice. On one hand, dynorphin acts postsynaptically via KORs on local GABAergic neurons in the VP to promote disinhibition of cholinergic neurons, which release acetylcholine into the amygdala to invigorate reward-seeking behaviors. On the other hand, dynorphin also acts presynaptically via KORs on dynorphinergic terminals to limit its own release. Such autoinhibition keeps cholinergic neurons from prolonged activation and release of acetylcholine, and prevents perseverant reward seeking. Our study reveals how dynorphin exquisitely modulate motivation through cholinergic system, and provides an explanation for why these neuromodulators are involved in motivational disorders, including depression and addiction.

## INTRODUCTION

Dynorphin is generated from the precursor protein prodynorphin (Pdyn) (Chavkin, 2013; Schwarzer, 2009). It acts primarily through κ-opioid receptors (KORs), a G_i/o_ type of G protein-coupled receptors (GPCRs) whose activation in general causes neuronal inhibition (Darcq and Kieffer, 2018). The inhibitory effects mediated by KORs have mostly been studied at presynaptic terminals, where activation of KORs causes suppression of neurotransmitter release. For example, in the nucleus accumbens (NAc), the bed nucleus of stria terminalis (BNST) and other brain areas, pharmacologic activation of KORs on presynaptic terminals of various types of inputs – dopaminergic, serotoninergic, GABAergic and glutamatergic – leads to inhibition of their release of the respective neurotransmitters (Ahrens et al., 2018; Pomrenze et al., 2022; Spanagel et al., 1992; Tejeda et al., 2017). Notably, the dynorphin/KOR signaling-mediated presynaptic inhibition, including that observed in the NAc and BNST, is typically accompanied by strong aversive or anxiogenic effects (Ahrens et al., 2018; Bruchas et al., 2011; Castro and Bruchas, 2019; Darcq and Kieffer, 2018; Donahue et al., 2015), which have led to the prevailing view that dynorphin and KORs form an “anti-reward” system in the brain (Darcq and Kieffer, 2018; Koob et al., 2014; Koob and Le Moal, 2008).

However, dynorphin/KOR signaling is also involved in reward-seeking behaviors through, at least in part, modulation of hypothalamic neurons (Cayir et al., 2024; Cooper et al., 1985; Morley and Levine, 1983; Sandoval-Caballero et al., 2024; Sandoval-Caballero et al., 2023), and recent studies show that stimulation of Pdyn neurons, which presumably induces dynorphin release, in the dorsal striatum (Xiao et al., 2020) or a subpopulation of Pdyn neurons in the NAc (Al-Hasani et al., 2015; Ibrahim et al., 2024) drives potent appetitive responses and positive reinforcement. These findings indicate that dynorphin/KOR signaling is more complicated than being simply anti-reward, and raise the possibility that the behavioral roles of this signaling may depend upon its actions in specific brain areas or the specific neural circuits therein.

Pdyn neurons in the NAc (NAc^Pdyn^ neurons) are the major source of dynorphin production in the brain (Castro and Bruchas, 2019). Apart from local release of dynorphin within the NAc, where it may exert its anti-reward function by, for example, presynaptic inhibition of dopaminergic inputs from the ventral tegmental area (Bals-Kubik et al., 1993; Spanagel et al., 1992) or glutamatergic inputs from the amygdala (Tejeda et al., 2017), in principle dynorphin should also be released by NAc^Pdyn^ neurons into downstream brain regions through their long-range projections. The ventral pallidum (VP; also known as substantia innominata) is the major output structure of the NAc (Heimer et al., 1997), and is a key structure involved in goal-directed motivation, including the motivation to pursue appetitive stimuli and the motivation to avoid aversive stimuli (Humphries and Prescott, 2010; Root et al., 2015; Smith et al., 2009; Stephenson-Jones et al., 2020; Wulff et al., 2019). In particular, recent studies demonstrate that different types of neurons in the VP – including cholinergic, GABAergic and glutamatergic neurons – have distinct roles in learning or invigorating valence-specific behaviors (Aitta-Aho et al., 2018; Crouse et al., 2020; Kimchi et al., 2024; Ottenheimer et al., 2020; Richard et al., 2016; Stephenson-Jones et al., 2020). Whether and how NAc^Pdyn^ neurons and their release of dynorphin participate in regulating VP neurons, thereby influencing motivated behaviors, remain unknown and warrant careful study.

Here, we investigated the roles of dynorphin/KOR signaling from NAc^Pdyn^→VP projections in regulating VP neuronal function and motivated behavior. Through molecular, genetic and optogenetic manipulations in combination with electrophysiology, *in vivo* real-time recording of dynorphin release in behaving animals and behavioral characterization, we uncover that dynorphin released from NAc^Pdyn^ neurons powerfully controls a disinhibitory circuit in the VP via both postsynaptic KORs on local GABAergic neurons and presynaptic KORs on NAc^Pdyn^ neuron axon terminals. Through this mechanism, dynorphin facilitates and furthermore finetunes the activation of VP cholinergic neurons and their release of acetylcholine (ACh) into the basolateral amygdala (BLA), which in turn invigorates reward-seeking behaviors.

## RESULTS

### NAc^Pdyn^ neurons are the major source of NAc outputs to the VP

Previous studies indicate that NAc^Pdyn^ neurons belong to dopamine receptor D1 (Drd1) type of medium spiny neurons (Al-Hasani et al., 2015; Ibrahim et al., 2024). Since Drd1 neurons are heterogenous (Xiao et al., 2020), we characterized them with single molecule fluorescent *in situ* hybridization (smFISH). These neurons are composed of two major populations, with one expressing *Pdyn* and the other expressing *Tshz1* (Supplementary Fig. 1A-D), akin to the Drd1 neurons in the dorsal striatum (Xiao et al., 2020). To examine the connectivity between these neurons and neurons in the VP, the major target of the NAc, we performed retrograde mono-transsynaptic tracing with rabies virus (RV), using VP neurons expressing either Gad2 (glutamate decarboxylase 2, a marker for GABAergic neurons) or ChAT (choline acetyltransferase, a marker for cholinergic neurons) as the starter cells (Supplementary Fig. 2A; Methods). We subsequently used smFISH to identify the types of RV-labeled neurons. For both classes of stater cells, most of the RV-labeled neurons in the NAc expressed *Pdyn* (Supplementary Fig. 2B-E; Gad2, 73%, ChAT, 82%). To verify the cell types directly innervated by NAc^Pdyn^ neurons, we performed anterograde mono-transsynaptic tracing with an HSV strain, using NAc^Pdyn^ neurons as the starter cells (Supplementary Fig. 2F). Subsequent smFISH revealed that the HSV-labeled neurons in the VP included both GABAergic neurons and cholinergic neurons, with GABAergic neurons being the majority (Supplementary Fig. 2G-K). These results indicate that NAc^Pdyn^ neurons provide the major source of NAc outputs to different neuronal types in the VP.

### NAc^Pdyn^ neurons disinhibit VP cholinergic neurons in a KOR-dependent manner

To examine the functional impact of NAc^Pdyn^ neuron outputs on VP neurons, we used *Pdyn^Cre^;ChAT^FlpO^* mice in which we injected the NAc and VP, respectively, with an adeno-associated virus (AAV) expressing the light-gated cation channel ChR2 in a Cre-dependent manner (AAV-DIO-ChR2) and an AAV expressing a red fluorescent protein mCherry in a Flp-dependent manner (AAV-fDIO-mCherry) (Fig. 1A). This strategy allowed selective expression of ChR2 in NAc^Pdyn^ neurons and mCherry in VP ChaT-expressing (VP^ChaT^) and thus cholinergic neurons. Acute slices containing the VP were prepared from these mice. We photo-stimulated axon fibers originating from NAc^Pdyn^ neurons, and recorded the evoked inhibitory postsynaptic currents (eIPSCs) from mCherry-positive (mCherry^+^) VP^ChaT^ neurons as well as adjacent mCherry-negative (mCherry^−^) neurons in the same slices (Fig. 1B). The latter population should mostly be GABAergic neurons as they are the predominate neuron type in the VP (Stephenson-Jones et al., 2020). Notably, VP^ChaT^ neurons exhibited much smaller GABA_A_-mediated eIPSCs and lower probability of exhibiting such eIPSCs than the putative GABAergic neurons (Fig. 1B, C). These results show that NAc^Pdyn^ neurons make stronger inhibitory synapses and have higher synaptic connectivity with GABAergic neurons than cholinergic neurons in the VP, and suggest that NAc^Pdyn^ neurons may drive disinhibition of the cholinergic neurons.

**Figure 1.**
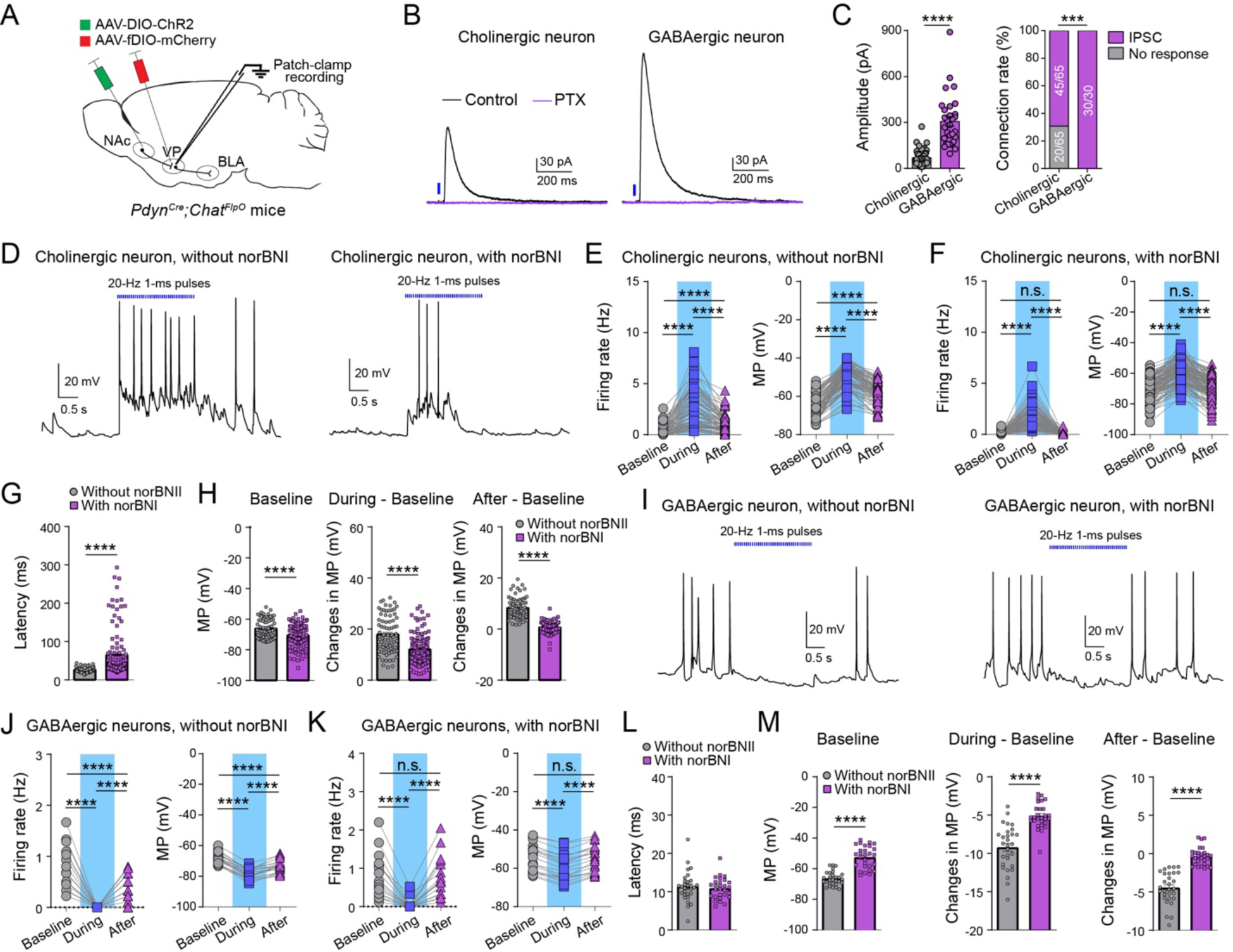
NAc^Pdyn^ neurons drive disinhibition of VP^ChAT^ neurons in a dynorphin/KOR signaling- dependent manner. (A) A schematic of the approach. (B) Traces of IPSCs recorded from a cholinergic neuron (left) and an adjacent putative GABAergic neuron (right) in the VP in the same slice in response to photo-stimulation of axon terminals originating from NAc^Pdyn^ neurons. (C) Quantification of IPSC amplitude (left) and connection rate (right) of cholinergic (mCherry^+^) and putative GABAergic (mCherry^−^) neurons (data was obtained from 7 mice; amplitude: cholinergic, n = 45/65 neurons, GABAergic, n = 30/30 neurons, Mann-Whitney test, ****P < 0. 0001; connection rate, cholinergic, 45 out of 65 neurons had measurable IPSCs, GABAergic, 30 out of 30 neurons had measurable IPSCs, Chi- square test, ***P = 0.0006). (D) Traces of current-clamp recording from two cholinergic neurons in response to photo-stimulation of axon terminals originating from NAc^Pdyn^ neurons. Left: without norBNI; right: with norBNI (100 nM). (E) Quantification of firing rate (left) and membrane potential (MP, right) before, during and after the photo- stimulation in the absence of norBNI (n = 79 neurons from 7 mice; firing rate, ****P < 0.0001; MP, ****P < 0.0001; Wilcoxon signed-rank test). (F) Same as E except that recording was performed in the presence of norBNI (n = 117 neurons from 7 mice; firing rate, ****P < 0.0001; n.s., nonsignificant, P = 0.5345; MP, ****P < 0.0001, n.s., P = 0.5519; Wilcoxon signed-rank test). (G) Quantification of the latency of light-evoked firing (without norBNI, n = 79 neurons from 7 mice; with norBNI, n = 117 neurons from 7 mice; ****P < 0.0001, Mann-Whitney test). (H) Quantification of baseline MP (left), changes in MP from baseline to photo-stimulation period (middle), and changes in MP from baseline to after photo-stimulation period (right) (****P < 0.0001, Mann-Whitney test). (I) Traces of current-clamp recording from putative GABAergic neurons in response to photo-stimulation of axon terminals originating from NAc^Pdyn^ neurons. Left: without norBNI; right: with norBNI (100 nM). (J) Quantification of firing rate (left) and MP (right) before, during and after the photo-stimulation in the absence of norBNI (n = 29 neurons from 3 mice; firing rate, ****P < 0.0001; MP, ****P < 0.0001; Wilcoxon signed-rank test). (K) Same as J except that recording was performed in the presence of norBNI (n = 28 neurons from 3 mice; firing rate, ****P < 0.0001; n.s., P = 0.2126; MP, ****P < 0.0001, n.s., P = 0.0726; Wilcoxon signed-rank test). (L) Quantification of the latency of light-evoked firing (without norBNI, n = 29 neurons from 3 mice; with norBNI, n = 28 neurons from 3 mice; P = 0.7097, Mann-Whitney test). (M) Quantification of baseline MP (left), changes in MP from baseline to photo-stimulation period (middle), and changes in MP from baseline to after photo-stimulation period (right) (without norBNI, n = 29 neurons from 7 mice; with norBNI, n = 28 neurons from 7 mice; ****P < 0.0001, Mann-Whitney test). Data are presented as mean ± s.e.m.

To test this possibility, we recorded the firing and membrane potential of VP^ChAT^ neurons in response to optogenetic stimulation of NAc^Pdyn^ terminals. Strikingly, the stimulation induced a marked increase in neuronal firing and depolarization, which lasted for seconds after cessation of the photo-stimulation (Fig. 1D-F). We reasoned that the lasting disinhibition is mediated by dynorphin signaling, which is slower than GABA_A_-mediated fast synaptic transmission. Indeed, application of KOR antagonist norbinaltorphimine (norBNI) decreased the resting membrane potential, decreased the light-evoked firing and depolarization, increased the latency of the firing, and completely blocked the lasting disinhibition of VP^ChAT^ neurons (Fig. 1D-H). These results suggest that dynorphin signaling makes an important contribution to the disinhibition of VP cholinergic neurons, including the slower, lasting component of the disinhibition.

Correspondingly, optogenetic stimulation of NAc^Pdyn^ terminals led to a dramatic decrease in firing and hyperpolarization of the putative VP GABAergic neurons (Fig. 1I-K), which lasted beyond the duration of photo-stimulation. These inhibitory effects were diminished by norBNI (Fig. 1I-M). These results together suggest that NAc^Pdyn^ neurons drive disinhibition of VP cholinergic neurons by inhibiting local GABAergic neurons, in a manner that depends on dynorphin/KOR signaling.

### KORs on GABAergic neurons are required for the disinhibition

To verify whether KORs on local GABAergic neurons contribute to the disinhibition of VP^ChAT^ neurons, we repeated the above experiments in *Gad2^Cre^* mice in which *Oprk1* expression in VP GABAergic (VP^Gad2^) neurons were selectively suppressed using CRISPR/Cas9-mediated mutagenesis. To this end, we used our recently developed AAV that conditionally expresses SaCas9 and a single-guide RNA targeting *Oprk1* (AAV1-FLEX-SaCas9-U6-sgOprk1) (Fellinger et al., 2021). An AAV with the same design but expressing a guide RNA that targets Rosa26 (AAV1-FLEX-SaCas9-U6-sgRosa26) was used as a control (Fellinger et al., 2021). We confirmed that expressing sg*Oprk1* in VP^Gad2^ neurons led to effective suppression of *Oprk1* (Supplementary Fig. 3). For the electrophysiology experiment, we used *Gad2^Cre^;ChAT^FlpO^*mice and injected the VP with the AAV that expresses sgOprk1, or the control sgRosa26, in a Cre-dependent (and thus GABAergic neuron-specific) manner, together with the AAV that expresses mCherry in a Flp-dependent (and thus cholinergic neuron-specific) manner (Supplementary Fig. 4A). In addition, we injected the NAc of the same mice with the Cre-dependent AAV to express ChR2 in GABAergic neurons, including NAc^Pdyn^ neurons.

We obtained whole-cell patch-clamp recording from mCherry^+^ VP^ChAT^ neurons in acute slices. In the sgRosa26 control group, photo-stimulation (2s, 20 Hz) of ChR2^+^ fibers originating from the NAc successfully induced disinhibition of VP^ChAT^ neurons, which lasted beyond the secession of light (Supplementary Fig. 4B-D). Administration of nor-BNI substantially reduced the disinhibition during photo-stimulation, and completely abolished the disinhibition after the secession of light. In contrast, in the sgOprk1 group, the photo-stimulation-induced disinhibition of VP^ChAT^ neurons was much reduced and did not last beyond the secession of light (Supplementary Fig. 4E-G). Application of norBNI did not further diminish the disinhibition, suggesting an occlusion effect (Supplementary Fig. 4H, I). These results indicate that KORs on local GABAergic neurons play a critical role in mediating the disinhibition of VP cholinergic neurons driven by NAc inputs.

### NAc^Pdyn^ neurons control reward-induced acetylcholine release in the BLA

To investigate the *in vivo* function of NAc^Pdyn^ neuron-driven disinhibition of VP cholinergic neurons, we virally expressed ChR2 in NAc^Pdyn^ neurons and the ACh sensor gACh3.0 (Jing et al., 2020) in the BLA (Fig. 2A), one of the major targets of VP cholinergic neurons (Crouse et al., 2020; Jiang et al., 2016). Optical fibers were implanted above the NAc and the BLA for photo- stimulation and photometry, respectively (Supplementary Fig. 5A, B). Remarkably, a single pulse of blue light delivered to the NAc with a duration as short as 50 ms was sufficient to trigger a robust gACh3.0 response in the BLA (Fig. 2B). Furthermore, photo-activation of the axon fibers in the VP originating from NAc^Pdyn^ neurons similarly triggered gACh3.0 response in the BLA (Fig. 2C, D; Supplementary Fig. 5C, D). In contrast, optogenetically activating Tshz1 or Drd2 (dopamine receptor D2) neurons in the NAc (targeted with *Tshz1^Cre^* or *Adora2a-Cre* mice, respectively) (Supplementary Fig. 6A-H), or activating the projections from hypothalamic Pdyn neurons to the VP (Supplementary Fig. 6I-L), failed to induce any gACh3.0 response in the BLA. These results suggest that NAc^Pdyn^ neurons have a specific role in driving disinhibition of VP cholinergic neurons, leading to ACh release in the BLA.

**Figure 2.**
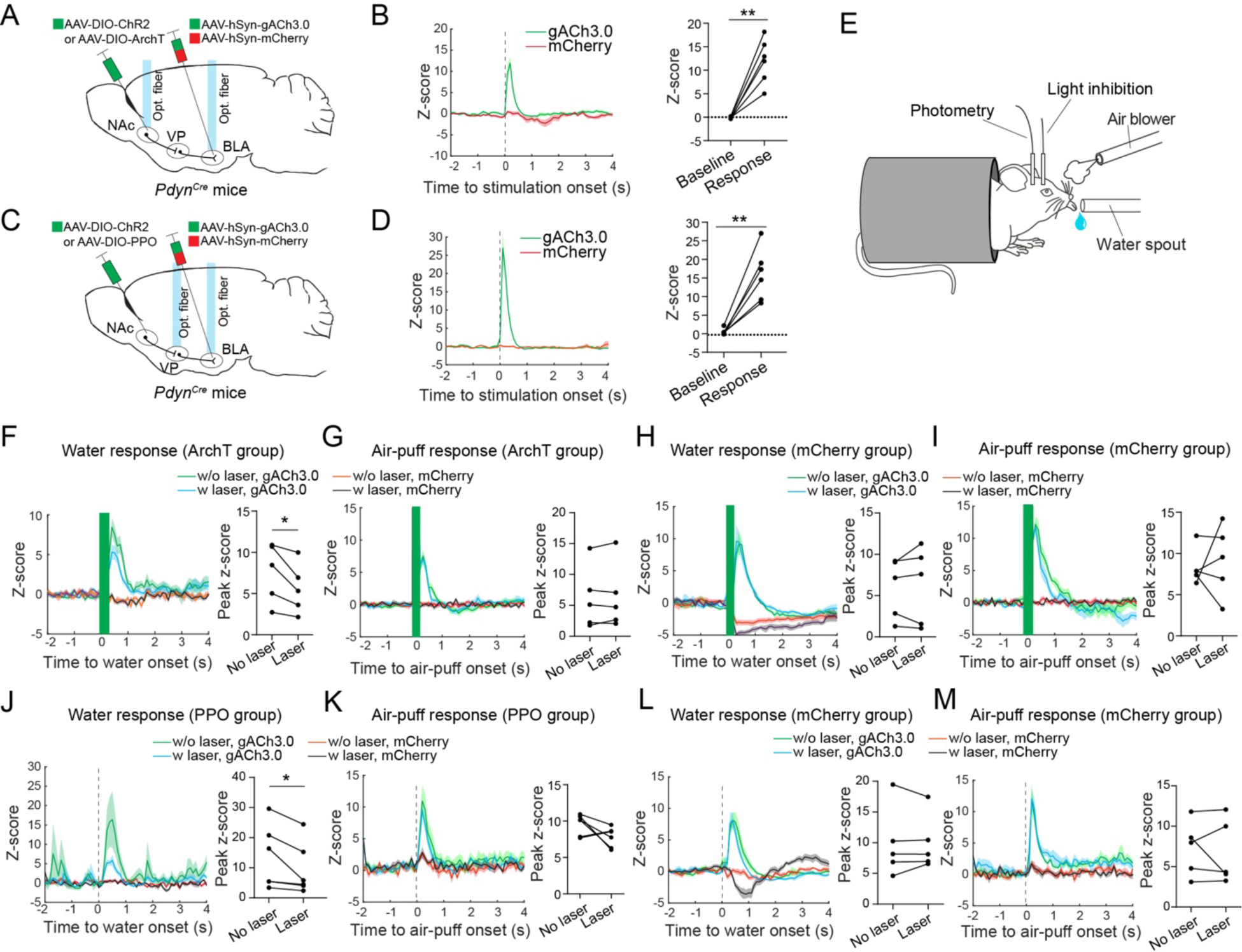
NAc^Pdyn^®VP projections control ACh release in the BLA. (A) A schematic of the approach. (B) Left: average gACh3.0 signals from a mouse that received photo-stimulation of ChR2-expressing Pdyn neurons in the NAc. mCherry signals are also shown to monitor potential motion artifacts. Right: quantification of the photo-stimulation-evoked response for all mice (n = 6 mice, t = 6.34, **P = 0.0014, paired t test). (C) A schematic of the approach. (D) Left: average gACh3.0 signals from a mouse that received photo-stimulation of axon terminals in the VP originating from ChR2-expressing NAc^Pdyn^ neurons. mCherry signals are also shown to monitor potential motion artifacts. Right: quantification of the photo-stimulation-evoked response for all mice (n = 6 mice, t = 5.86, **P = 0.0021, paired t test). (E) A schematic of the experimental design. (F) Left: average gACh3.0 signals from an ArchT mouse that received water. mCherry signals are also shown to monitor potential motion artifacts. Right: quantification of the response to water for all mice (n = 5 mice, t = 3.09, *P = 0.0366, paired t test). (G) Same as F, except that the mice received air-puff and the response was to air-puff (n = 5 mice, t = 0.56, P = 0.6, paired t test). (H) Left: average gACh3.0 signals from a mCherry mouse that received water. mCherry signals are also shown to monitor potential motion artifacts. Right: quantification of the response to water for all mice (n = 5 mice, t = 0.54, P = 0.62, paired t test). (I) Same as H, except that the mice received air-puff and the response was to air-puff (n = 5 mice, t = 0.42, P = 0.69, paired t test). (J) Left: average gACh3.0 signals from a PPO mouse that received water. mCherry signals are also shown to monitor potential motion artifacts. Right: quantification of the response to water for all mice (n = 5 mice, t = 2.68, *P = 0.044, paired t test). (K) Same as J, except that the mice received air-puff and the response was to air-puff (n = 5 mice, t = 1.9, P = 0.12, paired t test). (L) Left: average gACh3.0 signals from a mCherry mouse that received water. mCherry signals are also shown to monitor potential motion artifacts. Right: quantification of the response to water for all mice (n = 5 mice, t = 0.57, P = 0.96, paired t test). (M) Same as L, except that the mice received air-puff and the response was to air-puff (n = 5 mice, t = 0.47, P = 0.66, paired t test). Data are presented as mean ± s.e.m.

Next, we tested whether NAc^Pdyn^ neurons are required for ACh release in the BLA. We expressed the light-sensitive proton pump archaerhodopsin (ArchT) or mCherry (as a control) in NAc^Pdyn^ neurons, and expressed gACh3.0 in BLA neurons. Optical fibers were implanted above the infected areas in the NAc and BLA for light-inhibition and photometry, respectively (Fig. 2A). The mice were water restricted and presented with two stimuli that were either rewarding or aversive: water and air-puff blowing to the face, respectively (Fig. 2E). In randomly interleaved trials, we delivered brief (150 ms) green light pulses to the NAc during the water or air-puff presentations. Both water and air-puff triggered robust gACh3.0 response in the BLA (Fig. 2F, G). Interestingly, the water-evoked response, but not the air-puff-evoked response, in the ArchT mice was consistently reduced by the light pulses (Fig. 2F, G). In the mCherry mice, neither response was affected by the light pulses (Fig. 2H, I). These results suggest that inhibition of NAc^Pdyn^ neurons selectively impairs reward-evoked ACh release in the BLA.

To determine whether this effect is mediated by the NAc→VP pathway, we expressed parapinopsin (PPO), an opsin that causes rapid and sustained inhibition of presynaptic release upon blue light illumination (Copits et al., 2021), in NAc^Pdyn^ neurons (Fig. 2C). The mice were presented with water and air-puff as described above. In interleaved trials, we delivered blue light (10 mW, 10 s) into the VP to inhibit the NAc^Pdyn^ terminals before water or air-puff presentations (Fig. 2J-M; Methods). The light delivery substantially decreased gACh3.0 response to water, but did not affect that to air-puff (Fig. 2J, K). In mCherry control mice, light delivery into the VP had no effect on either the water-evoked or the air-puff-evoked gACh3.0 response (Fig. 2L, M). Together, these results indicate that NAc^Pdyn^ neurons specifically control reward-induced ACh release in the BLA through the NAc→VP pathway.

### NAc^Pdyn^**→**VP projections control reward-seeking behavior

To examine the behavioral role of the NAc→VP pathway, we sought to optogenetically manipulate NAc^Pdyn^→VP projections in mice during motivated behaviors. We first infected NAc^Pdyn^ neurons bilaterally with the AAV expressing PPO or mCherry and implanted optical fibers above the VP for light delivery (Fig. 3A, B). These mice were trained in a go/no-go task where two different tones (the conditioned stimuli, CS) predicted the delivery of either water or air-puff (the unconditioned stimuli, US) (Fig. 3B, C; Methods). During training, a 2-s pulse of blue light covering the time window between CS onset and US onset was delivered into the VP. Strikingly, compared with the control group, the PPO group had much reduced anticipatory licking in both the go and the no-go trials, resulting in reduced hit rate but increased correct rejection rate (CR) throughout training (Fig. 3D-G). The overall performance of the PPO group was impaired, mainly because of the large reduction in hit rate (Fig. 3G). These results suggest that inhibition of NAc^Pdyn^ axon terminals in the VP leads to impaired reward-seeking response in the go/no-go task.

**Figure 3.**
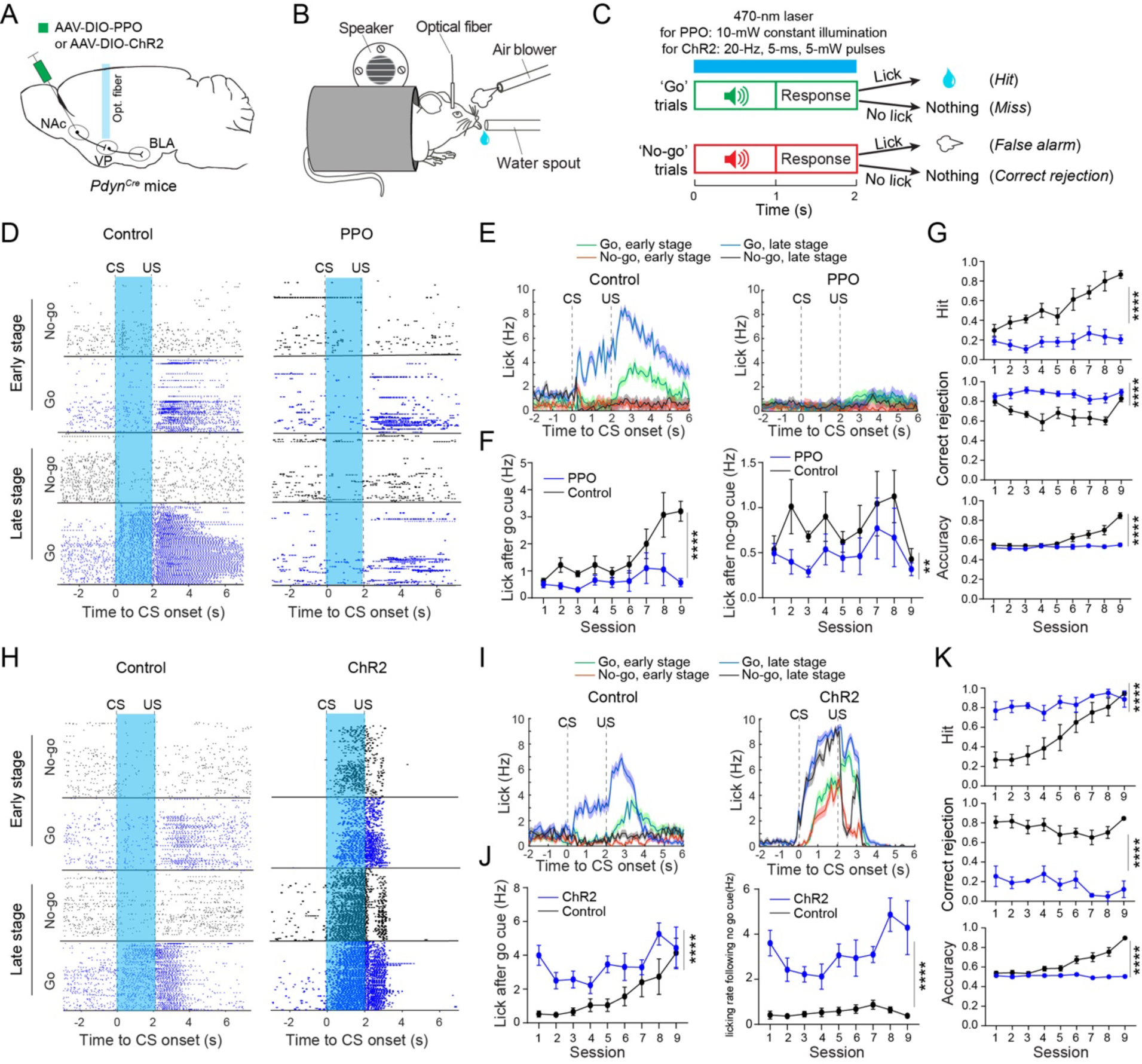
NAc^Pdyn^®VP projections control reward-seeking behavior. (A-C) Schematics of the approach (A), experimental setup (B) and task design (C). (D) Lick raster of a mCherry control mouse (left) and a PPO mouse (right) during go/no-go training. The blue shaded area indicates the time window when the laser was turned on. (E) Average licking rates of the same mice in D. (F) Left: quantification of licking rates following CS onset in go trials across training sessions (F(1,81) = 31.49, ****P = 2.7×10^-7^). Right: licking rates following CS onset in no-go trials across training sessions (F(1,81) = 8.679, **P = 0.0042). PPO group, n = 6 mice, mCherry group, n = 5 mice, two-way ANOVA followed by Sidak’s test. (G) Top: hit rate, F(1,81) = 142, ****P < 1×10^-15^. Middle: CR rate, F(1,81) = 74.96, ****P = 3.77×10^-13^. Bottom: accuracy, F(1,81) = 71.78, ****P = 8.78×10^-13^. PPO group, n = 6 mice, mCherry group, n = 5 mice, two-way ANOVA followed by Sidak’s test. (H) Lick raster of a mCherry mouse (left) and a ChR2 mouse (right) during go/no-go training. The blue shaded area indicates the time window when the laser was turned on. (I) Average licking rates of the same mice in H. (J) Left: quantification of licking rates following CS onset in go trials across training sessions (F(1,81) = 35.25, ****P = 7×10^-8^). Right: licking rate following CS onset in no-go trials across training sessions (F(1,81) = 110.4, ****P < 1×10^-15^). ChR2 group, n = 6 mice, mCherry group, n = 5 mice, two-way ANOVA followed by Sidak’s test. (K) Top: hit rate, F(1,81) = 67.66, ****P = 2.7×10^-12^. Middle: CR rate, F(1,81) = 332.3, ****P < 1×10^-15^. Bottom: accuracy, F(1,81) = 189.1, ****P < 1×10^-15^. ChR2 group, n = 6 mice, mCherry group, n = 5 mice. Two-way ANOVA followed by Sidak’s test. Data are presented as mean ± s.e.m.

We next infected NAc^Pdyn^ neurons bilaterally with the AAV expressing ChR2 or mCherry and implanted optical fibers above the VP for light delivery (Fig. 3A, B). These mice were also trained in the go/no-go task, in which a train of blue light pulses covering the time window between CS onset and US onset was delivered into the VP (Fig. 3B, C; Methods). Compared with the control group, the ChR2 group had much enhanced anticipatory licking in both the go and the no-go trials, resulting in increased hit rate but reduced CR throughout training (Fig. 3H- K). The overall performance of the ChR2 group was impaired, mainly because of the large reduction in CR (Fig. 3K). Thus, the effects of ChR2 are essentially opposite to those of PPO, and suggest that activation of the NAc^Pdyn^→VP projections leads to enhanced reward-seeking response in the go/no-go task.

### Reward induces dynorphin release in the VP

Our results from *ex vivo* brain slices suggest that dynorphin/KO signaling makes an important contribution to the disinhibition of VP cholinergic neurons driven by NAc^Pdyn^→VP projections (Fig. 1; Supplementary Fig. 4), while those from *in vivo* optogenetics suggest that these projections control reward-seeking behavior (Fig. 3). However, how endogenous dynorphin participates in modulating VP function and animal behavior is poorly understood, as *in vivo* real-time measurement of dynorphin dynamics has been infeasible until recently (Dong et al., 2024). To address this issue, we infected VP neurons with our recently developed genetically encoded dynorphin sensor kLight1.3 (Dong et al., 2024) and implanted an optical fiber above the infected area for recording kLight1.3 signals with photometry (Fig. 4A). Again, the mice were presented with water and air-puff, as described above (Fig. 2E). Notably, water evoked a clear increase in kLight1.3 signals, whereas air-puff caused a decrease in the signals (Fig. 4B, C).

**Figure 4.**
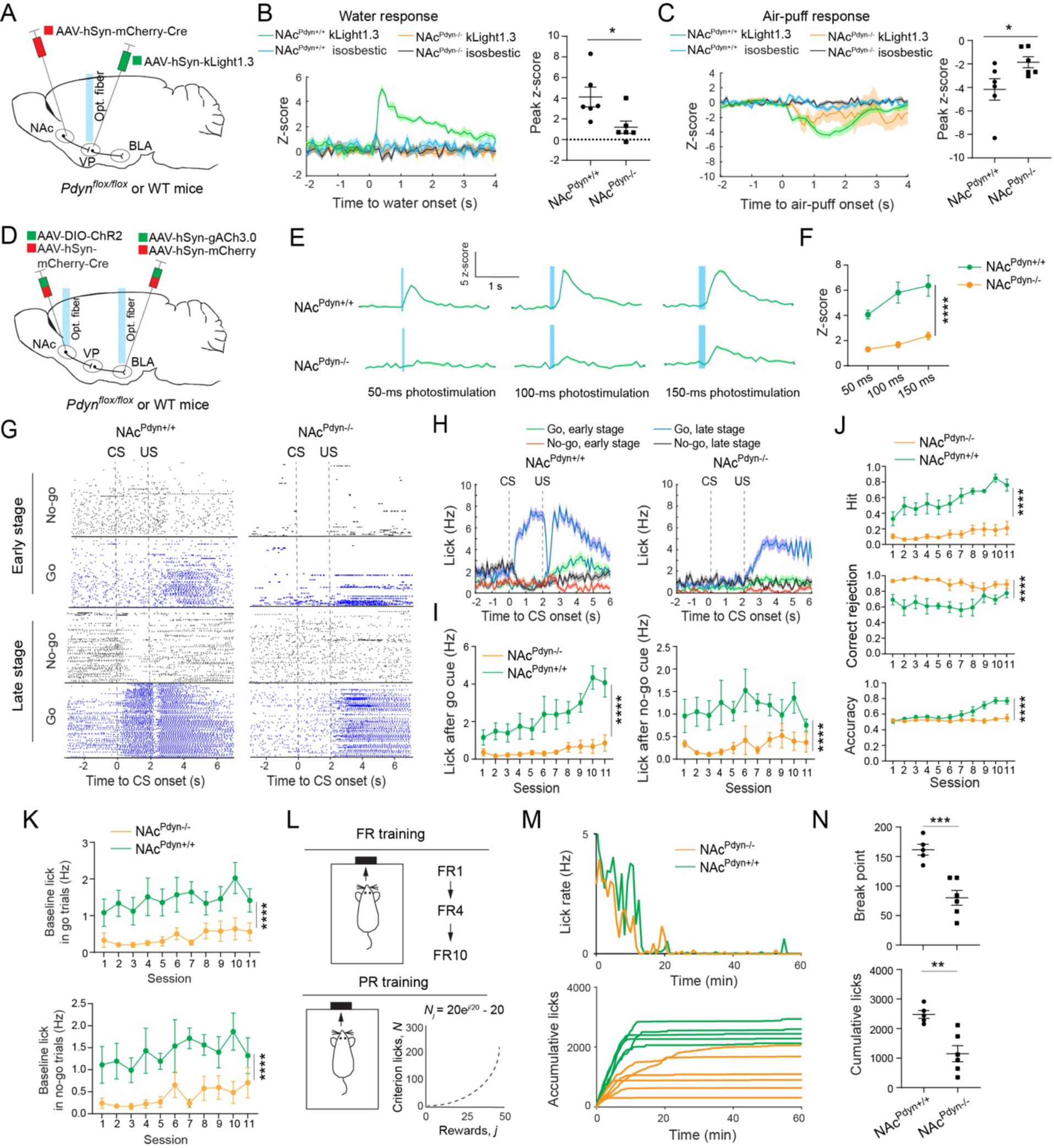
Dynorphin release into the VP is required for acetylcholine release in the BLA and motivation in mice. (A) A schematic of the approach. (B) Left, average kLight1.3 signals in the VP of a NAc^Pdyn+/+^ mouse and a NAc^Pdyn-/-^ mouse that received water. Signals from the isosbestic channel are also shown to monitor potential motion artifacts. Right, quantification of the response to water for all mice (n = 6 mice in each group, t = 2.59, *P = 0.027, unpaired t test). (C) Left, average kLight1.3 signals in the VP of a NAc^Pdyn+/+^ mouse and a NAc^Pdyn-/-^ mouse that received air-puff. Signals from the isosbestic channel are also shown to monitor potential motion artifacts. Right, quantification of the response to air-puff for all mice (n = 6 mice in each group, t = 2.25, *P = 0.0478, unpaired t test). (D) A schematic of the approach. (E) Average gACh3.0 signals in the BLA of a NAc^Pdyn+/+^ mouse (upper) and a NAc^Pdyn-/-^ mouse (lower) that received photo-stimulation of NAc neurons with different durations of light pulses. (F) Quantification of the photo-stimulation triggered response for all animals (n = 6 mice in each group, F(1,30) = 70.78, ****P = 2.17×10^-9^, two-way ANOVA followed by Sidak’s test). (G) Lick raster of a NAc^Pdyn+/+^ (control) mouse (left) and a NAc^Pdyn-/-^ mouse (right) during go/no-go training. (H) Average licking rates of the same mice in G. (I) Left: quantification of licking rates following CS onset in go trials across training sessions (F(1,99) = 104.8, ****P < 1×10^-15^). Right: licking rates following CS onset in no-go trials across training sessions (F(1,99) = 60.12, ****P = 8.06×10^-13^). NAc^Pdyn-/-^ group, n = 6 mice, NAc^Pdyn+/+^ group, n = 5 mice, two- way ANOVA followed by Sidak’s test. (J) Top: hit rate, F(1,99) = 235.3, ****P < 1×10^-15^. Middle: CR rate, F(1,99) = 100.5, ****P < 1×10^-15^. Bottom: accuracy, F(1,99) = 55.52, ****P = 3.52×10^-11^. NAc^Pdyn-/-^ group, n = 6 mice, NAc^Pdyn+/+^ group, n = 5 mice, two-way ANOVA followed by Sidak’s test. (K) Baseline licking rates during go/no-go training (upper: go trials, F(1,99) = 74.72, ****P = 9.8×10^-14^; lower: no-go trials, F(1,99) = 63.62, ****P = 2.7×10^-12^; two-way ANOVA followed by Sidak’s test). (L) Schematics of the design for fixed-ratio (FR) task and progressive-ratio (PR) task. (M) Upper: lick rate for a NAc^Pdyn-/-^ and NAc^Pdyn+/+^ mouse during PR test. Lower: cumulative licks for the NAc^Pdyn-/-^ and NAc^Pdyn+/+^ mice during PR test. (N) Quantification of break points (upper) and cumulative licks (lower) during PR test (break points, t = 5.05, ***P = 0.0007; cumulative licks, t = 4.11, **P = 0.0026; NAc^Pdyn-/-^ group, n = 6 mice, NAc^Pdyn+/+^ group, n = 5 mice, unpaired t test). Data are presented as mean ± s.e.m.

We repeated the above experiments in *Pdyn* conditional knockout (*Pdyn^flox/flox^*) mice (Bloodgood et al., 2021) in which *Pdyn* was deleted in NAc neurons by an AAV expressing Cre (NAc^Pdyn-/-^ mice; Fig. 4A-C). The deletion of *Pdyn*, which was confirmed with smFISH (Supplementary Fig. 7), markedly decreased water- or air-puff-evoked kLight1.3 signals in the VP of these mice as compared with the signals from their wild-type control (NAc^Pdyn+/+^) mice, indicating that the kLight1.3 signals represent dynorphin released by NAc^Pdyn^ neurons. These results demonstrate that a naturally rewarding stimulus triggers endogenous dynorphin release from NAc^Pdyn^ neurons into the VP.

### Dynorphin modulates ACh release in the BLA and motivation in mice

To determine the *in vivo* function of dynorphin released by NAc^Pdyn^ neurons, we first asked whether it is required for ACh release in the BLA. We deleted *Pdyn* in NAc neurons as described above and simultaneously expressed ChR2 in these neurons (Fig. 4D; Supplementary Fig. 7). The ACh sensor gACh3.0 was expressed in the BLA of the same mice. Brief light pulses delivered to the NAc only triggered modest gACh3.0 responses in the BLA of these NAc^Pdyn-/-^ mice, which were much smaller than the responses in NAc^Pdyn+/+^ mice (Fig. 4E, F). This result suggests that dynorphin released from NAc^Pdyn^ neurons is needed for effective ACh release in the BLA To verify this result and determine whether dynorphin acts via VP neurons, we pharmacologically blocked KOR with norBNI applied either systemically (Supplementary Fig. 8A, B; Methods) or locally within the VP (Supplementary Fig. 9A; Methods). *Pdyn^Cre^*mice were used in these experiments in which we measured ACh sensor gACh3.0 responses in the BLA to optogenetic activation of NAc^Pdyn^ neurons. In both cases, norBNI reduced the responses (Supplementary Fig. 8C, D; Supplementary Fig. 9B). Interestingly, systemic or local norBNI application also reduced gACh3.0 responses induced by water reward, but did not affect the responses induced by air-puff (Supplementary Fig. 8E, F; Supplementary Fig. 9C, D). In control animals in which saline was administered in lieu of norBNI, gACh3.0 responses in the BLA were stable (Supplementary Fig. 8G, H). These results together indicate that dynorphin released from NAc^Pdyn^ neurons is required for reward-induced ACh release in the BLA, likely because dynorphin/KOR signaling is critical for the disinhibition of VP cholinergic neurons (see Fig. 1; Supplementary Fig. 4).

To examine whether dynorphin released by NAc^Pdyn^ neurons is important for reward-driven behavior, we trained NAc^Pdyn-/-^ and control mice in the go/no-go task (Fig. 4G-J). The NAc^Pdyn-/-^ mice showed markedly lower anticipatory licking response and lower hit rate than the control mice in go trials (Fig. 4G-I). The NAc^Pdyn-/-^ mice also showed lower anticipatory licking response and therefore higher CR rate in no-go trials. Because of the marked reduction in hit rate, the overall performance of these mice was worse than the control mice (Fig. 4J). Notably, the NAc^Pdyn-/-^ mice showed much reduced baseline licking activity in both go and no-go trials (Fig. 4K). Consistent with these observations, the mice in which KOR was blocked by norBNI systemically or locally within the VP showed similar phenotypes (Supplementary Fig. 8I-N; Supplementary Fig. 9E-I). Of note, the effect of norBNI is long lasting (Ahrens et al., 2018; Al-Hasani et al., 2015; Chavkin et al., 2019).

The profound reduction in licking in these NAc^Pdyn-/-^ mice throughout the go/no-go task is reminiscent of the effects of PPO inhibition of the NAc^Pdyn^→VP projections (Fig. 3D-G), and prompted us to further examine whether these mice had reduced motivational drive using a progressive ratio (PR) task, whereby the effort required for thirsty animals to attain water increased progressively, and the breakpoint at which the mice stopped responding was used as a measure of motivation (Deng et al., 2021) (Fig. 4L). Indeed, the NAc^Pdyn-/-^ mice showed substantially decreased breakpoint and cumulative licking in the PR task (Fig. 4M, N), suggesting reduced motivation.

Recent studies show that stimulation of Pdyn neurons in the NAc can convey valence information and drive place preference in a real-time place preference (RTPP) test (Al-Hasani et al., 2015; Ibrahim et al., 2024). Therefore, we examined whether manipulations of dynorphin/KOR signaling in the NAc→VP pathway would affect behavior in the RTPP test. Activating NAc^Pdyn^ neurons or their projections to the VP induced robust RTPP (see all the control groups in Supplementary Fig. 10), consistent with the previous studies. Interestingly, activating NAc neurons sill induced robust RTPP in the NAc^Pdyn-/-^ mice (Supplementary Fig. 10A-C), or in the mice in which norBNI was administered systemically (Supplementary Fig. 10D-F) or locally within the VP (Supplementary Fig. 10G-I), suggesting that valence information from NAc^Pdyn^ neurons is conveyed by GABA rather than dynorphin. Together, these results suggest that dynorphin released from NAc^Pdyn^→VP circuit contributes to the disinhibition of VP cholinergic neurons and their subsequent ACh release into the BLA, and also modulates motivational drive thereby influencing reward-seeking responses in mice.

### KORs on VP GABAergic neurons are required for ACh release and motivation

We reasoned that KORs on VP GABAergic neurons are involved in controlling ACh release in the BLA and motivation in mice. To test this, we again used the AAV1-FLEX-SaCas9-U6- sg*Oprk1* to suppress *Oprk1* in VP GABAergic neurons of *Gad2^Cre^* mice (see Supplementary Fig. 3). In the BLA of the same mice, we expressed gACh3.0 and implanted optical fibers for recording ACh release with photometry (Fig. 5A). Similar to treatment with norBNI (Supplementary Fig. 8E-H; Supplementary Fig. 9C, D), suppressing *Oprk1* in VP GABAergic neurons reduced BLA gACh3.0 responses to water, but did not affect the responses to air-puff (Fig. 5B, C). Suppressing *Oprk1* in VP GABAergic neurons also reduced anticipatory licking and hit rate (Fig. 5D-G), as well as baseline licking (Fig. 5H) in the go/no-go task, and furthermore decreased breakpoint and cumulative licks in the PR test (Fig. 5I, J) in these animals, suggesting that they had reduced motivation. On the other hand, these mice behaved similarly to control animals in the elevated plus maze test (Supplementary Fig. 11), suggesting that suppressing *Oprk1* in VP GABAergic neurons does not affect basal anxiety levels.

**Figure 5.**
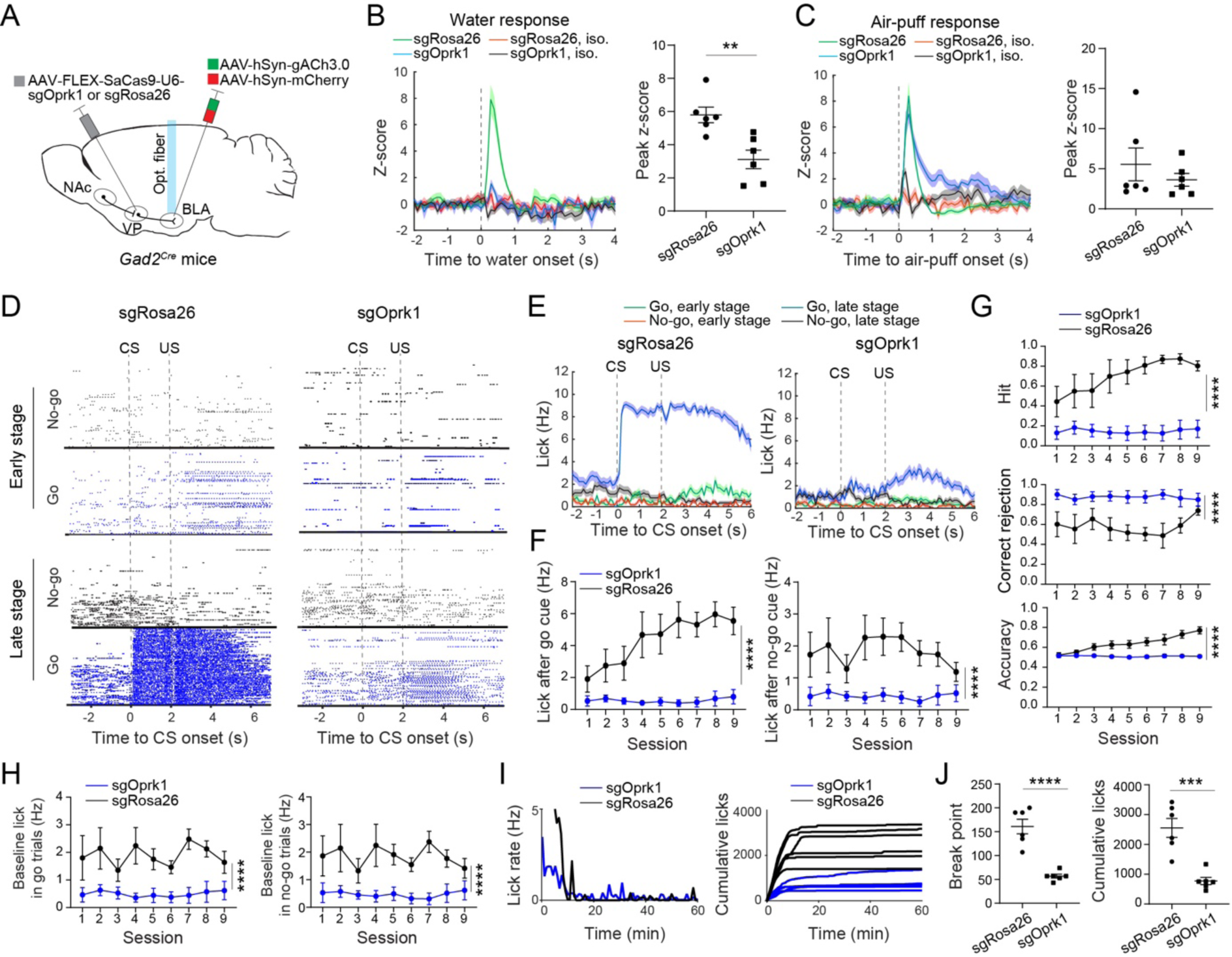
KORs on VP GABAergic neurons are required for acetylcholine release in the BLA and motivation in mice. (A) A schematic of the approach. (B) Left, average gACh3.0 signals in the BLA of a sgRosa26 (control) mouse and a sgOprk1mouse that received water. Signals from the isosbestic (iso.) channel are also shown to monitor potential motion artifacts. Right, quantification of the response to water for all mice (n = 6 mice in each group, t = 3.67, **P = 0.0043, unpaired t test). (C) Average gACh3.0 signals in the BLA of a sgRosa26 (control) mouse and a sgOprk1mouse that received air-puff. Signals from the isosbestic (iso.) channel are also shown to monitor potential motion artifacts. Right, quantification of the response to air-puff for all animals (n = 6 mice in each group, t = 0.88, P = 0.4, unpaired t test). (D) Lick raster of a sgRosa26 mouse (left) and a sgOprk1 mouse (right) during go/no-go training. (E) Average licking rates of the same mice in D. (F) Left: quantification of licking rates following CS onset in go trials across training sessions (F(1,90) = 113.1, ****P < 1×10^-15^. Right: licking rates following CS onset in no-go trials across training sessions (F(1,90) = 48.94, ****P = 4.53×10^-10^. N = 6 animals in each group, two-way ANOVA followed by Sidak’s test. (G) Top: hit rate, F(1,90) = 136.3, ****P < 1×10^-15^. Middle: CR rate, F(1, 90) = 57.15, ****P = 3.26×10^-11^. Bottom: accuracy, F(1,90) = 105.6, ****P < 1×10^-15^. N = 6 animals in each group, two-way ANOVA followed by Sidak’s test. (H) Baseline licking rates during go/no-go training (left: go trials, F(1,90) = 52.6, ****P = 1.37×10^-10^; right: no-go trials, F(1,90) = 47.56, ****P = 7.16×10^-10^; two-way ANOVA followed by Sidak’s test). (I) Left: lick rate for a sgRosa26 and sgOprk1 mouse during PR test. Right: cumulative licks for the sgRosa26 and sgOprk1 mice during PR test. (J) Quantification of break points (left) and cumulative licks (right) during PR test (break points, t = 6.53, ****P = 0.00007; cumulative licks, t = 5.22, ***P = 0.0004; sgOprk1 group, n = 6 mice, sgRosa26 group, n = 6 mice, unpaired t test). Data are presented as mean ± s.e.m.

To examine whether MORs on VP GABAergic neurons are required for valence processing, we suppressed *Oprk1* in these neurons as above, and expressed ChR2 in GABAergic neurons in the NAc, where optical fibers were also implanted for photo-stimulation. These mice were subjected to the RTPP test. We found that these mice showed robust preference for the stimulation side, similar to control mice (Supplementary Fig. 10J-L). This result is consistent with those from the NAc^Pdyn-/-^ mice (Supplementary Fig. 10A-C) and mice in which norBNI was administered systemically (Supplementary Fig. 10D-F) or locally within the VP (Supplementary Fig. 10G-I), confirming that dynorphin/KOR signaling in the NAc→VP pathway is not essential for valence processing. Rather, our results together suggest that this signaling is critical for motivational control, likely by modulating VP cholinergic neurons and their release of ACh in the BLA.

### The autoinhibitory function of KORs on NAc^Pdyn^ neurons

Previous studies show that KORs are highly expressed in Drd1 neurons in the NAc (Tejeda et al., 2017). Indeed, we found that *Oprk1* is expressed in almost all NAc^Pdyn^ neurons (Supplementary Fig. 12A, B), which is the major subtype of NAc Drd1 neurons (Supplementary Fig. 1). Since KORs can mediate presynaptic inhibition of inhibitory synapses (Ahrens et al., 2018; Tejeda et al., 2017), we reasoned that KORs on NAc^Pdyn^ neurons may gate the output of these neurons via autoinhibition. To test this, we injected the NAc of *Pdyn^Cre^* mice with the sgOprk1 (or the control sgRosa26) virus together with a ChR2 virus to infect NAc^Pdyn^ neurons. In the BLA of the same mice, we expressed gACh3.0. Optical fibers were implanted in the NAc for photo-stimulation and the BLA for photometry (Fig. 6A). The sgOprk1 virus led to suppression of *Oprk1* in NAc^Pdyn^ neurons (Supplementary Fig. 12A, C), as expected.

**Figure 6.**
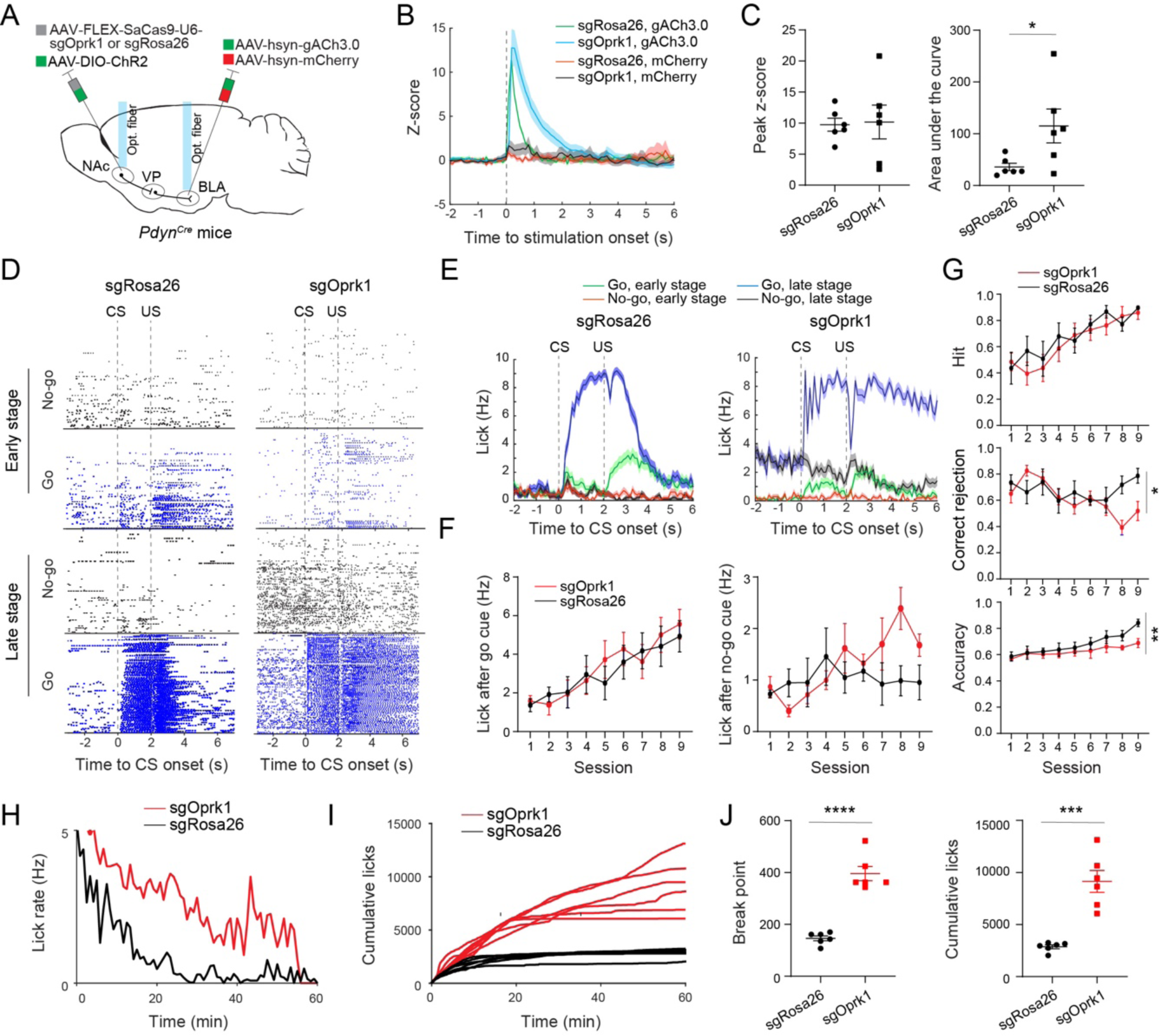
KORs on NAc^Pdyn^ neurons limit acetylcholine release in the BLA and motivation in mice. (A) A schematic of the approach. (B) Average gACh3.0 signals in the BLA of a sgRosa26 (control) mouse and a sgOprk1mouse that received photo-stimulation in the NAc. mCherry signals are used to monitor potential motion artifacts. (C) Quantification of the photo-stimulation-evoked response for all mice. Left: amplitude, t = 0.18, P = 0.88; right: area under the curve, t = 2.37, *P = 0.039; n = 6 mice for each group, unpaired t test. (D) Lick raster of a sgRosa26 mouse (left) and a sgOprk1 mouse (right) during go/no-go training. (E) Average licking rates of the same mice in D. (F) Left: quantification of licking rates following CS onset in go trials across training sessions, F(1,90) = 0.3026, P = 0.5836. Right: licking rates following CS onset in no-go trials across training sessions, F(1,90) = 3.247, P = 0.075. N = 6 animals in each group, two-way ANOVA followed by Sidak’s test. (G) Top: hit rate, F(1,90) = 1.021, P = 0.315. Middle: correct rejection rate, F(1,90) = 3.968, *P = 0.049. Bottom: accuracy, F(1,90) = 8.83, **P = 0.0038. N = 6 animals in each group, two-way ANOVA followed by Sidak’s test. (H) Lick rate for a sgRosa26 and sgOprk1 mouse during PR test. (I) Cumulative licks for the sgRosa26 and sgOprk1 mice during PR test. (J) Quantification of break points (left) and cumulative licks (right) during PR test (break points, t = 8.54, ****P = 6.58×10^-6^; cumulative licks, t = 5.88, ***P = 0.0002; n = 6 mice in each group, unpaired t test). Data are presented as mean ± s.e.m.

Remarkably, the photo-stimulation-triggered gACh3.0 responses in the sgOprk1 mice lasted much longer than those in the control mice, although the response amplitude was similar between the two groups (Fig. 6B, C). Consistently, in acute slices, the NAc^Pdyn^-neuron-driven disinhibition of VP^ChAT^ neurons in the sgOprk1 group lasted longer than the control group (Supplementary Fig. 13). These results indicate that a normal function of KORs on NAc^Pdyn^ neurons is to gate the output of these neurons via autoinhibition, thereby limiting the disinhibition of VP^ChAT^ neurons and their release of ACh in the BLA.

To determine how KORs on NAc^Pdyn^ neurons might influence motivated behavior, we suppressed *Oprk1* in NAc^Pdyn^ neurons with the sgOprk1 virus, and trained these animals and their controls in the go/no-go task (Fig. 6D-F). Compared to the control mice, the sgOprk1 mice showed enhanced licking in no-go trials (Fig. 6D-F), resulting in decreased CR rate and overall performance (Fig. 6G). In addition, in the PR test, the sgOprk1 mice showed markedly increased breakpoint and cumulative licking (Fig. 6H-J). On the other hand, photo-stimulation of NAc^Pdyn^ neurons where *Oprk1* was suppressed induced similar levels of RTPP to control mice (Supplementary Fig. 10M-O), suggesting normal valence processing. These results indicate that an impairment in dynorphin/KOR signaling in NAc^Pdyn^ neurons leads to abnormally enhanced motivational drive for reward, even when there is a punishment (such as the air-puff in the no-go trials, Fig. 6D-G).

### VP cholinergic output to the BLA is critical for motivational control

Our results thus far point to a possibility that VP cholinergic output to the BLA modulates motivation. To test this possibility, we first examined the effects of activating VP→BLA cholinergic projections. We expressed ChR2 (or YFP, as a control) in VP^ChAT^ neurons in *ChAT^FlpO^* mice, and implanted optical fibers in the BLA of the same mice for light delivery (Fig. 7A; Supplementary Fig. 14A-E). These mice were trained to perform the PR task, in which trains of blue light pulses were delivered into the BLA in interleaved trials (Methods). We found that the photo-stimulation consistently increased the breakpoint and cumulative licking in the ChR2 mice (Fig. 7B, C), while it had no effect on the control mice (Fig. 7D, E), indicating that activating VP→BLA cholinergic projections elevates animals’ motivational drive. Photo- stimulation of these projections did not affect animal behavior in the RTPP test (Supplementary Fig. 14F, G), suggesting that they do not convey valence information.

**Figure 7.**
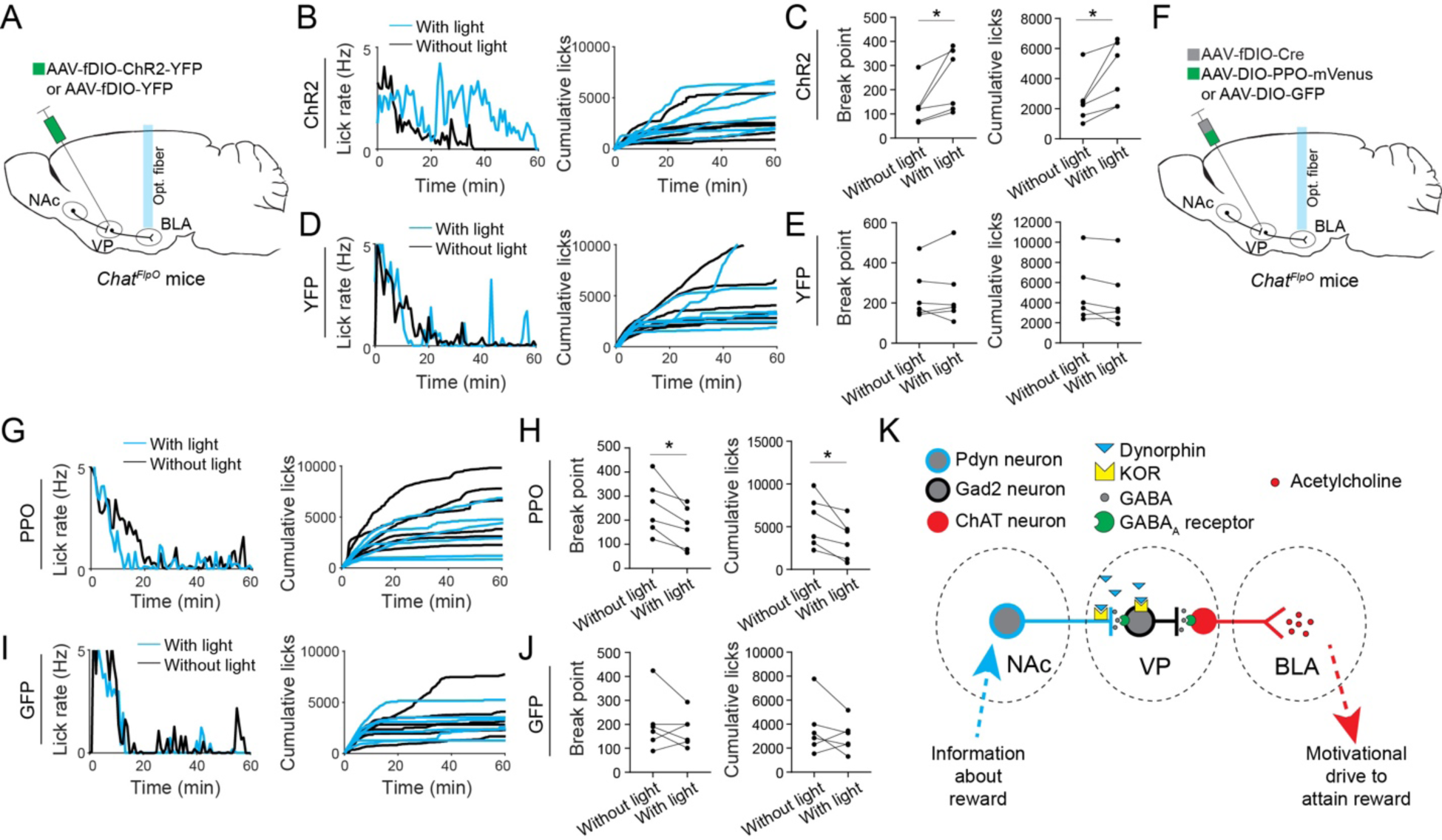
VP®BLA cholinergic projections control motivation. (A) A schematic of the approach. (B) Left: lick rate for a ChR2 mouse during PR test. Right: cumulative licks for the ChR2 mice during PR test. (C) Quantification of break points (left) and cumulative licks (right) for the ChR2 mice during PR test (break points, t = 2.7, *P = 0.0428; cumulative licks, t = 3.04, *P = 0.0289; n = 6 mice in each group, unpaired t test). (D) Left: lick rate for a YFP mouse during PR test. Right: cumulative licks for the YFP mice during PR test. (E) Quantification of break points (left) and cumulative licks (right) for the YFP mice during PR test (break points, t = 0.3, P = 0.78; cumulative licks, t = 1.7, P = 0.15; n = 6 mice in each group, unpaired t test). (F) A schematic of the approach. (G) Left: lick rate for a PPO mouse during PR test. Right: cumulative licks for the PPO mice during PR test. (H) Quantification of break points (left) and cumulative licks (right) for the PPO mice during PR test (break points, t = 3.86, *P = 0.119; cumulative licks, t = 3.2, *P = 0.0243; n = 6 mice in each group, unpaired t test). (I) Left: lick rate for a GFP mouse during PR test. Right: cumulative licks for the GFP mice during PR test. (J) Quantification of break points (left) and cumulative licks (right) for the GFP mice during PR test (break points, t = 0.84, P = 0.44; cumulative licks, t = 1.07, P = 0.33; n = 6 mice in each group, unpaired t test). (K) A circuit model of motivational control. Data are presented as mean ± s.e.m.

Finally, we examined the effects of inhibiting VP→BLA cholinergic projections. We expressed PPO (or GFP, as a control) in VP^ChAT^ neurons in *ChAT^FlpO^* mice, and implanted optical fibers in the BLA of the same mice (Fig. 7F; Supplementary Fig. 14H-J). These mice were also trained to perform the PR task, during which a constant blue light was delivered into the BLA (Methods). Interestingly, light delivery into the BLA led to a consistent decrease in breakpoint and cumulative licking in the PPO mice (Fig. 7G, H), but it did not affect the behavior of the control mice (Fig. 7I, J), indicating that inhibiting VP→BLA cholinergic terminals suppresses animals’ motivational drive. These results together indicate that the cholinergic inputs from the VP to the BLA bidirectionally modulate motivational drive.

## DISCUSSION

Our results support a model in which endogenous dynorphin released by NAc^Pdyn^ neurons in response to reward powerfully modulates a disinhibitory circuit in the VP, which controls activation of cholinergic neurons and their release of ACh into the BLA, thereby controlling the motivation to obtain reward (model Fig. 7K). In this disinhibitory circuit – which consists of the projections from NAc^Pdyn^ neurons to the VP, local VP GABAergic (VP^Gad2^) neurons and VP cholinergic (VP^ChAT^) neurons projecting to the BLA – activation of NAc^Pdyn^ neurons causes potent and lasting inhibition of VP^Gad2^ neurons and disinhibition of VP^ChAT^ neurons, leading to robust ACh release into the BLA. Each of these processes appears to be critical for reward seeking behavior. Notably, dynorphin release by NAc^Pdyn^ neurons, which occurs in response to reward delivery, plays an indispensable and intricate role in these processes. The released dynorphin acts through KORs in two locations within this circuit: VP^Gad2^ neurons and NAc^Pdyn^ neuron projections to the VP, which are postsynaptic and presynaptic, respectively, with respect to the site of dynorphin production (Fig. 7K). Through activation of KORs on VP^Gad2^ neurons, dynorphin enables potent and lasting inhibition of these neurons – an effect that outlasts GABA_A_ receptor-mediated fast synaptic inhibition – and thus disinhibition of VP^ChAT^ neurons and the subsequent release of ACh into the BLA. This function of dynorphin is ultimately required for the motivation of animals to pursue reward. By contrast, through activation of KORs on NAc^Pdyn^ neurons, dynorphin induces autoinhibition of their projections to the VP, which finetunes the disinhibition of VP^ChAT^ neurons and the release of ACh into the BLA, thereby adjusting motivational drive. Our study thus uncovers cellular and circuit mechanisms that are essential for motivational control.

A notable observation, which is enabled by the newly developed dynorphin sensor (Dong et al., 2024), is that dynorphin release from NAc^Pdyn^ neurons into the VP is triggered by a rewarding stimulus, but not by an aversive one. This observation is at first glance surprising, as it is at odds with the prevailing notion that dynorphin/KOR conveys anti-reward signals. However, it is consistent with our further observation that dynorphin invigorates reward-seeking behaviors through the VP^ChAT^→BLA cholinergic circuit, and also consistent with previous reports that Pdyn neurons (Al-Hasani et al., 2015; Ibrahim et al., 2024; Xiao et al., 2020) or dynorphin/KOR signaling (Cooper et al., 1985; Morley and Levine, 1983; Sandoval-Caballero et al., 2024; Sandoval-Caballero et al., 2023) are involved in reward-seeking behaviors. The notion that dynorphin/KOR signaling is “anti-reward” is likely rooted in the specific brain areas it is examined and the methods used for the examination. For instance, infusion of KOR agonists into the NAc can result in the suppression of transmitter release from various inputs, including dopaminergic inputs and glutamatergic inputs that are critical for reward processing (Castro and Bruchas, 2019; Darcq and Kieffer, 2018), therefore causing aversive or dysphoric effects. Our study, on the other hand, probes the functions of dynorphin and KORs in the distinct elements of VP circuits, which are known to control motivation during reward-seeking behaviors (Aitta-Aho et al., 2018; Crouse et al., 2020; Kimchi et al., 2024; Ottenheimer et al., 2020; Richard et al., 2016; Stephenson-Jones et al., 2020). Thus, the seemingly contradictory findings about dynorphin/KOR function can be explained by the divergent behavioral roles of neural circuits whereby this signaling system acts.

Neurons in subcortical nuclei, such as the NAc, the bed nucleus of the stria terminalis, the central amygdala and hypothalamic areas, often express various neuropeptides that can be co-released with fast-acting neurotransmitters from the same neurons (Castro and Bruchas, 2019; Fellinger et al., 2021; Li et al., 2022; Soden et al., 2023). The development of CRISPR/Cas9 mutagenesis viruses has made it possible to study their functions in isolation (Hunker et al., 2020). The present study is in line with recent studies showing that neuropeptides have functions distinct from those of fast-acting neurotransmitters (Li et al., 2022; Soden et al., 2023). In our study, deletion of dynorphin or KORs in the NAc^Pdyn^→VP^Gad2^ circuit does not affect behavior in the RTPP test, suggesting that the slow-acting dynorphinergic transmission in this circuit is not essential for valence processing. Therefore, it will be interesting to further examine whether valence processing is dependent on fast-acting neurotransmitters in this circuit.

The cholinergic system has been implicated in multiple brain functions, including attention, learning, mood regulation, reward processing and motivation (Collins et al., 2016; Mineur and Picciotto, 2021, 2023; Nunes et al., 2022). In particular, cholinergic neurons in the VP send dense projections to the BLA, which contribute to both fear learning and reward learning (Aitta-Aho et al., 2018; Crouse et al., 2020; Jiang et al., 2016; Woolf, 1991; Zaborszky et al., 2012) and promote reward responding (Kimchi et al., 2024). Our results extend these findings, showing that these projections are both sufficient and necessary for the motivation to pursue reward. Interestingly, it has been shown that BLA projections to the NAc drive reward-seeking behaviors (Britt et al., 2012; Namburi et al., 2015; Stuber et al., 2011). Thus, a BLA→NAc^Pdyn^→VP^Gad2^→VP^ChAT^→BLA loop may exist and be critical for maintaining the motivation for reward seeking. Alternatively, or in addition, projections from BLA Fezf2 neurons to the olfactory tubercle have recently been shown to drive reward-seeking behavior (Zhang et al., 2021) and thus may also mediate ACh function in motivational control. Future studies will delineate how ACh regulates distinct BLA circuits to influence motivation as well as other brain functions. Moreover, future studies also need to examine whether alterations in the NAc^Pdyn^→VP^Gad2^→VP^ChAT^→BLA circuit contribute to motivational disorders, including depression and drug addiction.

## ACKNOWLEDGEMENTS

We thank Richard Palmiter for permission to use the *Pdyn^flox/flox^* mouse line, Karl Deisseroth for generously providing the AAV5-EF1α-fDIO-hChR2(H134R)-YFP virus, Radhashree Sharma, Charlotte Lee and Darren Chen for technical assistance, and members of the Li laboratory for helpful discussions. This work was supported by grants from National Institutes of Health (NIH) (R01MH108924, R01DA050374, R01MH101214, R01NS104944, B.L.), the Cold Spring Harbor Laboratory and Northwell Health Affiliation (B.L.), and the Key R&D Program of Zhejiang (2024SSYS0031).

## AUTHOR CONTRIBUTIONS

Q.S., M.L., W.G. and B.L. designed research; Q.S., M.L., and W.G. performed research; Q.S., M.L., W.G. and B.L. analyzed data; X.X. assisted with the PR task; C.D. and L.T. developed the dynorphin sensor; M.R.B. assisted with the *Pdyn^flox/flox^*mouse line; L.S.Z. developed the CRISPR/Cas9 system targeting *Oprk1*; Y.L. developed the acetylcholine sensor; B.L. supervised the study; and Q.S. and B.L. wrote the paper with inputs from all authors.

## RESOURCE AVAILABILITY

### Lead Contact

Further information and requests for resources and reagents should be directed to and will be fulfilled by the Lead Contact, Bo Li.

### Materials Availability

This study did not generate new unique reagents.

### Data and code availability

- The authors declare that the data supporting the findings of this study are available within the paper and its supplementary information files.
- All original code has been deposited at Figshare and is publicly available as of the date of publication.
- Any additional information required to reanalyze the data reported in this paper is available from the lead contact upon request.

## EXPERIMENTAL MODEL AND SUBJECT DETAILS

Male and female mice (2–4 months old) were used for all the experiments. Mice were housed under a 12-h light/12-h dark cycle (light from 7 a.m. to 7 p.m.) with a constant room temperature of 21 °C and 65% humidity. Mice were housed in groups of 2–5. Food and water were available ad libitum before the start of experiments. All experiments were performed during the light cycle. Littermates were randomly assigned to different groups before the experiments. All experimental procedures were approved by the Institutional Animal Care and Use Committee of Cold Spring Harbor Laboratory and performed in accordance with the US National Institutes of Health guidelines.

*Pdyn^Cre^* (Strain #:027958), *Gad2^Cre^*(Strain #:010802), *Chat^Cre^* (Strain #:031661), *Chat^FlpO^* (Strain #:036281), and C57BL/6J (Strain #:000664) mice were purchased from the Jackson Laboratory. *Adora2a-Cre* mice (RRID: MMRRC_036158-UCD) were purchased from MMRRC. The *Pdyn^flox/flox^* mouse line was described previously (Bloodgood et al., 2021).

The *Tshz1^Cre^* knock-in mouse driver line, in which the expression of Cre recombinase is driven by the endogenous *Tshz1* promoter, was generated as previously described (Xiao et al., 2020). A gene-targeting vector for *Tshz1^Cre^* was generated using a PCR-based cloning approach (Taniguchi et al., 2011) to insert a *2A-Cre* construct immediately before the STOP codon of the *Tshz1* gene. The targeting vector was linearized and transfected into a 129SVj/B6 F1 hybrid ES cell line (V6.5, Open Biosystems). G418-resistant ES clones were first screened by PCR and then confirmed by Southern blotting using probes against the 5’ and 3’ homology arms of the targeted site.

## METHOD DETAILS

### Viral vectors

AAV-EF1a-DIO-hChR2(H134R)-EYFP-WPRE-HGHpA (AAV5, 1.38×10^13^ genome copies (GC) per ml), AAV-hSyn-DIO-mCherry (AAV2, 2×10^13^ GC per ml), AAV-EF1a-fDIO-mCherry (AAV5, 2.3×10^13^ GC per ml), AAV-hSyn-mCherry (AAV8, 2.6×10^13^ GC per ml), AAV-Ef1a- DIO-PPO-mVenus (AAV9, 1×10^13^ GC per ml), AAV-EF1a-fDIO-Cre (AAV8, 1×10^13^ GC per ml) were purchased from Addgene. rAAV9/CAG-FLEX-ArchT-GFP (4.7 × 10^12^ GC per ml), AAV8-hSyn-mCherry-Cre (5 × 10^12^ GC per ml) were purchased from University of North Carolina Vector Core Facility. AAV-hSyn-gACh3.0 (1.3× 10^13^ GC per ml) was purchased from WZ bio. AAV-hSyn-klight1.3 was generated by Lin Tian lab. AAV1-Flex-SaCas9-sgOprk1, AAV1-Flex-SaCas9-U6-sg-Rosa26 and AAV1-Flex-eGFP-Kash was generated by Larry Zweifel lab. AAV5-EF1α-fDIO-hChR2(H134R)-YFP (5 × 10^12^ GC per ml) was generated by Deisseroth lab. AAV9-CAGGS-Flex-mKate-T2A-TVA (5 × 10^12^ GC per ml), AAV9- CAGGS-Flex-mKate-T2A-N2c-G (5 × 10^12^ GC per ml) and Rbv-CVS-N2c-dG-GFP (5 × 10^8^ plaque-forming units (PFUs) per ml) were generated by HHMI Janelia Research Campus. AAV-DIO-EGFP-2A-TK (2.66 × 10^12^ GC per ml) and HSV-dTK-hUbc-tdTomato (1.0 × 10^9^ PFUs per ml) were purchased from BrainVTA. All viral vectors were aliquoted and stored at −80 °C until use.

### Stereotaxic surgery and virus injection

All surgeries were performed under aseptic conditions, and body temperature was maintained with a heating pad. Standard surgical procedures were used for stereotaxic injection and implantation, as previously described (Xiao et al., 2020; Yang et al., 2023; Zhang et al., 2021). Briefly, mice were anesthetized with isoflurane (2% at the beginning for induction and 1–1.5% for the rest of the surgery) and positioned in a stereotaxic frame. The frame was linked to a digital mouse brain atlas to guide the targeting of different brain structures (Angle Two Stereotaxic System, Leica Biosystems Division of Leica). The following stereotaxic coordinates in anteroposterior axis (AP), mediolateral axis (ML), and dorsoventral axis (DV) (all in mm in reference to Bregma) were used for the NAc: AP 1.1, ML 1.2, and DV -4.8; for the VP: AP 0, ML 1.5, and DV -5; for the BLA: AP -1.65, ML 3.3, and DV -4.8; and for the VMH: AP -1.6, ML 0.35, and DV -5.5. 200 – 300 nl of viral solution was injected at a speed of 1-2 nl/s. For AAVs, we typically waited at least 3-4 weeks after the injection to allow viral expression. For RV tracing, we injected the TVA and G helper viruses (at a ratio of 1:2 (volume:volume), 150 nl in total) first, and injected the RV (300 nl) 2-3 weeks later. We waited for 7 days after RV injection before collecting the brains for histological analysis. For HSV tracing, we injected the helper virus (150 – 200 nl) first, and injected the HSV (200 nl) 3 weeks later. We waited for 5 days after HSV injection before collecting the brains for histological analysis.

To examine the functional projections from the NAc to VP neurons and the disinhibition effects, AAV-EF1a-DIO-hChR2(H134R)-EYFP-WPRE-HGHpA was bilaterally injected into the NAc of *Pdyn^Cre^;ChAT^FlpO^*mice (7-8 weeks old) in a volume of 300 nl for each site, and AAV-EF1a- fDIO-mCherry was bilaterally injected into the VP of the same mice in a volume of 300 nl for each side. These mice were subjected to slice electrophysiology experiments 2-3 weeks later. To check the disinhibition of cholinergic neurons in vivo, 300 nl AAV-EF1a-DIO-hChR2(H134R)- EYFP-WPRE-HGHpA was bilaterally injected into the NAc of *Pdyn^Cre^* mice, and a mixture of AAV-hSyn-mCherry and AAV-hSyn-gACh3.0 (1:10 in volume) was injected into the BLA. For the optogenetic activation in Supplementary Fig. 6, 300 nl of AAV-EF1a-DIO-hChR2(H134R)- EYFP-WPRE-HGHpA was bilaterally injected into the NAc of *Tshz1^Cre^* or *Adora2a-Cre* mice, and a mixture of AAV-hSyn-mCherry and AAV-hSyn-gACh3.0 (1:10, volume:volume) was injected into the BLA. For the activation of Pdyn^+^ terminals from the hypothalamus, 300 nl of AAV-EF1a-DIO-hChR2(H134R)-EYFP-WPRE-HGHpA was bilaterally injected into the VMH of *Pdyn^Cre^* animals, and a mixture of AAV-hSyn-mCherry and AAV-hSyn-gACh3.0 (1:10, volume:volume) was injected into the BLA. For optogenetic inhibition, 300 nl of AAV-Ef1a- DIO-PPO-Venus was bidirectionally injected into the NAc, and a mixture of AAV-hSyn-mCherry and AAV-hSyn-gACh3.0 (1:10, volume:volume) was injected into the BLA.

For measuring the dynorphin sensor klight1.3 with fiber photometry, 300 nl of AAV8-hSyn-mCherry-Cre was bidirectionally injected into the NAc of *Pdyn^flox/flox^* animals or wild-type (WT) control animals. 300 nl of AAV-hSyn-klight1.3 was injected into the VP. After 4 weeks of virus expression, the animals were used for fiber photometry. To measure acetylcholine release in the BLA in response to photo-simulation of NAc neurons with or without *Pdyn* expression in the NAc, a mixture of AAV8-hSyn-mCherry-Cre and AAV-EF1a-DIO-hChR2(H134R)-EYFP-WPRE-HGHpA (1:1, volume:volume) was injected into the NAc, and a mixture of AAV-hSyn-mCherry and AAV-hSyn-gACh3.0 (1:10, volume:volume) was injected into the BLA. After 4 weeks of virus expression, the animals were used for fiber photometry.

To test the role of KORs in GABAergic neurons in the VP, AAV1-Flex-SaCas9-sgOprk1 or AAV1-Flex-SaCas9-U6-sg-Rosa26 was bidirectionally injected into the VP of *GAD2^Cre^* mice, and a mixture of AAV-hSyn-mCherry and AAV-hSyn-gACh3.0 (1:10, volume:volume) was injected into the BLA. After 4 weeks of virus expression, the animals were used for fiber photometry. For the RTPP test in mice in which the expression of KORs was knocked down in GABAergic neurons in the VP, AAV1-Flex-SaCas9-sgOprk1 or AAV1-Flex-SaCas9-U6-sg- Rosa26 was bidirectionally injected into the VP, and 300 nl of AAV-EF1a-DIO-hChR2(H134R)- EYFP-WPRE-HGHpA was bilaterally injected into the NAc of of *GAD2^Cre^*mice. To determine the role of KORs on VP GABAergic neurons in disinhibiting VP cholinergic neurons, AAV- EF1a-DIO-hChR2(H134R)-EYFP-WPRE-HGHpA was bilaterally injected into the NAc of *Gad2^Cre^;ChAT^FlpO^*mice (7-8 weeks old) in a volume of 300 nl for each site, and a mixture of AAV-EF1a-fDIO-mCherry and AAV1-Flex-SaCas9-sgOprk1 (or AAV1-Flex-SaCas9-U6-sg- Rosa26) was bilaterally injected into the VP in a volume of 300 nl for each side (1:1, volume:volume). These mice were subjected to slice electrophysiology experiments 2-3 weeks later.

To test the role of KORs in NAc^Pdyn^ neurons, a mixture of AAV-EF1a-DIO-hChR2(H134R)- EYFP-WPRE-HGHpA and AAV1-Flex-SaCas9-sgOprk1 (or AAV1-Flex-SaCas9-U6-sg-Rosa26) was bilaterally injected into the NAc of *Pdyn^Cre^* mice in a volume of 300 nl for each side (1:1, volume:volume). A mixture of AAV-hSyn-mCherry and AAV-hSyn-gACh3.0 (1:10, volume:volume) was injected into the BLA for fiber photometry. For electrophysiology recording in acute slices, a mixture of AAV-EF1a-DIO-hChR2(H134R)-EYFP-WPRE-HGHpA and AAV1-Flex-SaCas9-sgOprk1 (or AAV1-Flex-SaCas9-U6-sgRosa26) was injected into the NAc *Pdyn^Cre^;ChAT^FlpO^*mice in a volume of 300 nl (1:1, volume:volume). 300 nl of AAV-EF1a- fDIO-mCherry was injected into the VP of the same mice to label cholinergic neurons.

For photo-activation of cholinergic terminals in the BLA, AAV5-EF1α-fDIO-hChR2(H134R)- YFP was bilaterally injected into the VP of *ChAT^Flpo^* mice in a volume of 300 nl for each side. For photo-inhibition of cholinergic terminals in the BLA, a mixture of AAV-Ef1a-DIO-PPO-Venus and AAV-EF1a-fDIO-Cre was bilaterally injected into the VP of *ChAT^Flpo^*mice in a volume of 300 nl for each side (1:1, volume:volume).

For *in vivo* optogenetics, optical fiber implantation was performed after viral injection in the same surgery. Optical fibers (core diameter, 200 µm; length, 5 mm; NA, 0.22; Inper Corporation) were implanted bilaterally and placed 200 µm above the NAc or VP. For *in vivo* fiber photometry, optical fiber implantation was also performed after viral injection in the same surgery. Optical fibers (core diameter, 200 µm; length, 5 mm; NA, 0.37; Inper Corporation) were placed unilaterally in the BLA or VP. A metal head-bar (for head restraint in all mice used in the photometry and behavioral experiments) was subsequently mounted onto the skull with black dental cement. We waited for a minimum of 3-4 weeks before starting the behavioral experiments in these mice.

### Immunohistochemistry

Immunohistochemistry experiments were performed following standard procedures described previously (Deng et al., 2021; Xiao et al., 2020). Briefly, mice were anesthetized with Euthasol (0.2 ml; Virbac, Fort Worth, Texas, USA) and transcardially perfused with 30 ml PBS, followed by 30 ml 4% paraformaldehyde (PFA) in PBS. Brains were extracted and further fixed in 4% PFA overnight followed by cryoprotection in a 30% PBS-buffered sucrose solution for 36-48 h at 4 °C. Coronal sections (50 μm) were cut using a freezing microtome (Leica SM 2010R, Leica). Sections were first washed in PBS (5 min), incubated in PBST (0.3% Triton X-100 in PBS) for 30 min at room temperature (RT) and then washed with PBS (3 x 5 min). Next, sections were blocked in 5% normal goat serum in PBST for 30 min at RT and then incubated with primary antibodies overnight at 4 °C. Sections were washed with PBS (3 x 5 min) and incubated with fluorescent secondary antibodies at RT for 2 h. In some experiments (as indicated in Figures and Supplementary Figures), sections were washed twice in PBS, incubated with DAPI (4′,6- diamidino-2-phenylindole, Invitrogen, catalogue number D1306) (0.5µg/ml in PBS) for 2 min. After washing with PBS (3 x 5 min), sections were mounted onto slides with Fluoromount-G (eBioscience, San Diego, California, USA). Images were taken using an LSM 780 laser-scanning confocal microscope (Carl Zeiss, Oberkochen, Germany). The primary antibodies used were: chicken anti-GFP (Aves Labs, catalogue number GFP1020; dilution 1:1000), rabbit anti-RFP (Rockland, catalogue number 600-401-379; dilution 1:1000), Anti-choline acetyltransferase (ChAT) antibody (Sigma-Aldrich, catalogue number AB144P; dilution 1:1000). Appropriate fluorophore-conjugated secondary antibodies (Life Technologies) were used depending on the desired fluorescence colors.

### Fluorescence *in situ* hybridization

Single-molecule fluorescent *in situ* hybridization (smFISH) (RNAscope, ACDBio) was used to detect the expression of *Pdyn*, *mCherry*, *GFP*, *Oprk1*, *Gad2*, *Tshz1*, *Drd1*, *Drd2*, *tdTomato*, *Vglut2* and *ChAT*. For tissue preparation, mice were first anesthetized with isoflurane and then decapitated. Their brain tissue was first embedded in cryomolds (Sakura Finetek, Catalog number 4566) filled with M-1 Embedding Matrix (Thermo Scientific, Catalog number 1310) and then quickly fresh-frozen on dry ice. The tissue was stored at −80 °C until it was sectioned with a cryostat. Cryostat-cut sections (16-μm thick) containing the brain areas of interest were collected along the rostro- caudal axis in a series of four slides and quickly stored at −80 °C until being processed. Hybridization was carried out using RNAscope kit (ACDBio). On the day of the experiment, frozen sections were postfixed in 4% PFA in RNA-free PBS (hereafter referred to as PBS) at room temperature (RT) for 15 min, then washed in PBS, dehydrated using increasing concentrations of ethanol in water (50%, once; 70%, once; 100%, twice; 5 min each). Sections were then dried at RT and incubated with Protease IV for 30 min at RT. Sections were washed in PBS three times (5 min each) at RT and hybridized.

Probes against *Pdyn* (Catalog number 318771-C2, dilution 1:50), *mCherry* (Catalog number 431201-C3, dilution 1:50), *GFP* (Catalog number 400281-C2, dilution 1:50), *Oprk1* (Catalog number 316111-C1, dilution 1:50), *Gad2* (Catalog number 439371-C2 and C3, dilution 1:50), *Tshz1* (Catalog number 494291-C3, dilution 1:50), *Drd1* (Catalog number 406491-C1, dilution 1:50), *Drd2* (Catalog number 406501-C3, dilution 1:50), *slc17a6* (Catalog number 319171-C3, dilution 1:50), *tdTomato* (Catalog number 317041-C1, dilution 1:50), and *ChAT* (Catalog number 410071-C3, dilution 1:50) were applied to the brain sections. Hybridization was carried out for 2 h at 40 °C. After that, the sections were washed twice in 1× Wash Buffer (Catalog number 310091; 2 min each) at RT, and then incubated with the amplification reagents for three consecutive rounds (30 min, 15 min and 30 min, at 40 °C). After each amplification step, the sections were washed twice in 1× Wash Buffer (2 min each) at RT. Finally, fluorescence detection was carried out for 15 min at 40 °C. Sections were then washed twice in 1× Wash Buffer (2 min each), incubated with DAPI for 2 min, washed twice in 1× Wash Buffer (2 min each), and then mounted with a coverslip using mounting medium.

Images were acquired using an LSM780 confocal microscope equipped with 20x and 40x lenses and visualized and processed using ImageJ and Adobe Illustrator. Cell counting and mean fluorescence intensity quantification of images were performed using ImageJ. To compare gene deletion or knock-down efficiency, brain sections were imaged with the same imaging settings and the fluorescence intensities of targeted genes in the experimental groups were normalized to those of the control groups.

### *In vitro* electrophysiology

Patch-clamp recording was performed as described previously (Li et al., 2013; Zhang et al., 2021). Adult (8- to 16-week-old) mice were deeply anesthetized by an overdose of isoflurane. Their brains were extracted, and coronal brain slices (290 µm thick) were generated at a slicing speed of 0.12 mm/s in ice-cold cutting solution containing 110 mM choline chloride, 25 mM NaHCO_3_, 1.25 mM NaH_2_PO_4_, 2.5 mM KCl, 0.5 mM CaCl_2_, 7.0 mM MgCl_2_, 25.0 mM glucose, 11.6 mM ascorbic acid and 3.1 mM pyruvic acid (osmolarity, 300-310 mOsm), gassed with 95% O_2_ and 5% CO_2_ using a vibrating-blade microtome (HM650, Thermo Fisher Scientific). The slices were transferred to a holding chamber and incubated in 34 °C artificial cerebrospinal fluid (ACSF) containing: 118 mM NaCl, 2.5 mM KCl, 26.2 mM NaHCO_3_, 1 mM NaH_2_PO_4_, 20 mM glucose, 2 mM MgCl_2_, and 2 mM CaCl_2_ (osmolarity, 300-310 mM, pH, 7.4), which was oxygenated with 95% O_2_ and 5% CO_2_. Forty-five minutes after recovery, the slices were transferred to a recording chamber and perfused with oxygenated ACSF at 3 ml/min at room temperature (20-24 °C).

Whole-cell patch-clamp recordings were performed using glass pipettes with a resistance of 3-7 MΩ. A blue LED (470 nm, pE-100, CoolLED) was used to activate ChR2. The 470-nm light-evoked postsynaptic responses of VP cholinergic neurons were recorded in voltage-clamp mode. The internal solution contained 115 mM cesium methanesulfonate, 20 mM CsCl, 10 mM HEPES, 2.5 mM MgCl_2_, 4 mM Na_2_ATP, 0.4 mM Na_3_GTP, 10 mM sodium phosphocreatine and 0.6 mM EGTA (pH 7.2, osmolarity ∼295 mOsm). IPSCs were recorded at a voltage of 0 mV. CNQX (10 μM) and D-AP5 (50 μM) was added into the ACSF to block glutamate receptors. Picrotoxin (PTX; 50 μM) was used to block GABA_A_ receptors. For disinhibition-related experiments, cholinergic or GABAergic neurons in the VP were recorded in current-clamp mode (holding current, 0 pA) using a potassium-based internal solution containing (in mM) 130 K-gluconate, 5 KCl, 2.5 MgCl_2_, 10 HEPES, 0.6 EGTA, 10 mM sodium phosphocreatine 0.4 Na_3_-GTP, and 4 Na_2_-ATP (pH 7.2, osmolarity ∼290 mOsm). CNQX (10 μM) and D-AP5 (50 μM) was added into the ACSF to block excitatory synaptic inputs. In some experiments, norBNI (nor- Binaltorphimine Dihydrochloride, Sigma-Aldrich, catalogue number 5.08017, 100 nM) was added into the ACSF to block κ-opioid receptors. Action potential firing was obtained every 30 s with light stimulation (a 2-s train of 20-Hz 1-ms light pulses). The locations of VP cholinergic neurons were identified with an air objective (5X; NA 0.10).

Voltage clamp and current clamp recordings were carried out using a MultiClamp 700B amplifier (Molecular Devices). During recording, traces were low-pass filtered at 3 kHz (Digidata 1440A; Molecular Devices). Data were acquired with Axon Clampex 10.2 software. The amplitude of inhibitory postsynaptic currents (IPSCs) was analyzed using pCLAMP 10 and Igor software. IPSC amplitudes were calculated as the difference between the peak amplitude within 50 ms after light stimulation onset and the mean amplitude just before the IPSC. For recording of spontaneous membrane potential and firing rate, cells were held in current clamp mode and no current injections were made. Membrane potential and firing were recorded for at least 2 s before and after light stimulation. Action potential latency was measured as the time difference from the onset of the first light stimulation to the half-height of the peak of the first action potential.

### Measuring acetylcholine release in behaving mice

To measure acetylcholine release in response to water and air-puff, custom-built spouts were used to deliver these stimuli to mice. An external trigger from a Bpod State Machine (Sanworks) was used to synchronize the delivery with fiber photometry recording. A water-restriction schedule started 23 h before training. Mice were first habituated in a head-restraint frame for 10 min each day for 2 days. On day 3, mice were trained to lick water from the water spout. Once mice learned to successfully lick water, they were provided with water during fiber photometry recording. In randomly interleaved trials, air-puff was delivered toward the face of the mice. Each session contained 10 trials of water delivery and 10 trials of air-puff delivery. To inhibit NAc^Pdyn^ neurons optogenetically with ArchT and determine the effects of the inhibition on acetylcholine release, a constant green light (532 nm, 10 mW) was delivered into the NAc starting at 50 ms before stimulus (water or air-puff) onset and ending at 100 ms after stimulus (water or air-puff) was ended. Trials with or without the light were randomly interleaved, with inter-trial intervals of 3-7 seconds. There were 10 trials for each trial type (water without light, air-puff without light, water with light, and air-puff with light). To inhibit NAc^Pdyn^ neuron axion terminals optogenetically with PPO and determine the effects of the inhibition on acetylcholine release, a 10-s constant blue light (470 nm, 10 mW) was delivered into the VP during each inter-trial interval. At 2 s following the cessation of the light, a stimulus (water of air-puff) was delivered to the animal. Trials with or without the light were randomly interleaved, with inter-trial intervals of 21-25 seconds. There were 5 trials for each trial type (water without light, air-puff without light, water with light, and air-puff with light). To activate NAc^Pdyn^ neurons or their axion terminals optogenetically with ChR2 and determine the effects of the activation on acetylcholine release, an external trigger generated by a Bpod State Machine (Sanworks) was used to synchronize the delivery of blue light pulses (470 nm, 5 mW; 50, 100, or 150 ms in duration) into the NAc or VP with fiber photometry recording in the BLA. Each session consisted of 20 trials.

### In vivo fiber photometry and data analysis

We used a commercial fiber photometry system (FP3001, Neurophotometrics) to record signals from the dynorphin sensor (klight1.3) and the acetylcholine sensor (gACh3.0; see above “Measuring acetylcholine release in behaving mice”) *in vivo* in behaving animals under head restraint, through optical fibers (fiber core diameter, 200 µm; fiber length, 5.0 mm; NA, 0.37; Inper) implanted in the VP or BLA. A patch cord (fiber core diameter, 200 µm; Doric Lenses) was used to connect the photometry system with the implanted optical fibers. The intensity of the blue light (λ = 470 nm) for excitation was adjusted to a low level (20–50 µW) at the tip of the patch cord. Emitted sensor fluorescence was bandpass filtered and focused on the sensor of a CCD camera. Photometry signals (frame rate, 20 Hz) and behavioral events were aligned on the basis of an analogue TTL signal generated by the Bpod. Mean values of signals from a region of interest were calculated and saved using Bonsai software (Bonsai) and were exported to MATLAB (2017a) for further analysis. To correct for photobleaching of fluorescence signals (baseline drift), a bi-exponential curve was fitted to the raw fluorescence trace and subtracted as follows:

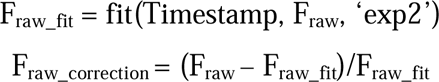

After baseline drift correction, the fluorescence signals were z-scored relative to the mean and standard deviation of the signals of the entire trace, excluding the time window when laser stimulation occurred (to avoid the light bleed-through from photo-stimulation). Besides the sensor signals, we simultaneously recorded isosbestic signals or mCherry signals which served to monitor potential motion artifacts. Trials with clear motion artifacts were excluded from further analysis.

### Go/no-go task

We trained mice in a go/no-go task as previously described (Xiao et al., 2020). Mice underwent a water-deprivation schedule that started 23 hours before training, and then 2 days of habituation to head restraint. After habituation, mice were trained to lick for water from a metal spout (5 µl per lick, 200 trials per session, 1 session per day for 3 days). Once mice learned to successfully obtain water in at least 85% of the trials, they were subjected to training in the go/no-go task (200 trials per session (100 go trials and 100 no-go trials), 1 session per day). In each trial, a 1-s pure tone cue (the CS) (go cue, 10 kHz, 70 dB; no-go cue, 3 kHz, 70 dB) was presented, followed by a 1-s delay. The delay was designated as the ‘decision window’. During go trials, the mice were required to lick at least once during the decision window to receive a drop of water (US; 10 µl), resulting in a hit trial. If mice did not lick during the decision window, they would not receive the water reward, resulting in a miss trial. During no-go trials, if mice licked the spout at least once during the decision window, they would receive a blow (200 ms) of air-puff, resulting in a ‘false alarm’ (FA) trial. If mice did not lick during the window, they would successfully prevent air-puff delivery, resulting in a ‘correct rejection’ (CR) trial. Training in this phase persisted until the mice reached a performance level of at least 80% successful trials. The accuracy was calculated as the total correct responses divided by the total trials: accuracy = (hits + correct rejects) / (total trials).

For optogenetic activation, blue light (λ = 470 nm; 5 mW; 20-Hz 5-ms pulses) was delivered during the CS period and decision window. For optogenetic inhibition with PPO, blue light (λ = 470 nm; 10 mW; a 2-s square pulse) was delivered during the CS period and decision window.

For training in the go/no-go task after systemic norBNI application, mice received norBNI (25mg/kg, dissolved in 0.9% saline solution) intraperitoneal (i.p.) injection 3 weeks after they had received virus injection and fiber implantation. The animals were subjected first to fiber photometry recording, both before and 24 hours after the norBNI injection, and subsequently to training in the go/no go task. For training in the go/no-go task after norBNI local application in the VP, mice received norBNI (2.5 mg/ml, 500 nl) injection into the VP 3 weeks after they had received virus injection. Optical fibers were implanted subsequently. One week later, the animals were subjected to fiber photometry measurement. They were subsequently subjected to training in the go/no go task. For training in the go/no-go task after deletion of *Pdyn* or *Oprk1*, mice were subjected to training in the go/no-go task 4 weeks after they had received injection of the virus to delete the respective gene.

### Progressive ratio task

The progressive ratio (PR) task was modified based on a similar task described in a previous study (Deng et al., 2021). Water restricted mice were placed in a head-fixed rig equipped with a water port. Mice were first trained to lick into the port for water reward on a ‘‘fixed ratio 1’’ (FR1) schedule for 2 days, during which every lick leads to a reward (3 µl of water). Following the FR1 training, the schedule was changed to FR4 for 2 days, which required the mice to lick 4 times with a maximal interlick-interval of 2 minutes in order to receive the reward. Next, the schedule was changed to FR10 for 1 day. Finally, mice were tested with a progressive ratio (PR) schedule in which the number of licks required to obtain one reward followed a geometric progression according to a function:

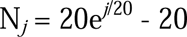

where j is the trial number. The function was modified on the basis of previous studies (Hodos, 1961; Richardson and Roberts, 1996). Before the PR schedule, mice were tested in an FR10 session. For the optogenetic activation during PR, mice received 20 Hz photo-stimulation (5-ms pulses; 4-s laser on periods with 2-s laser off intervals; power, 5 mW; λ = 473 nm) during the entire PR session (60 min). For the optogenetic inhibition during PR with PPO, mice received constant blue light (power, 10 mW, λ = 470 nm) during the entire PR session (60 min).

### Real-time place aversion or preference test

Freely moving mice were initially habituated to a two-sided chamber (23 cm × 33 cm × 25 cm; made from Plexiglas) for 10 min, during which their baseline preference for the left or right side of the chamber was assessed. The test consisted of two sessions (10 min each). During the first session, we assigned one side of the chamber (counterbalanced across mice) as the photo-stimulation side and placed the mice in the non-stimulation side to start the experiment. Once the mouse entered the stimulation side, photo-stimulation (5-ms pulses, 20 Hz, 5 mW measured at the tip of the optic fibers) generated by a 470-nm laser (OEM Laser Systems) was immediately turned on and was turned off as soon as the mouse exited the stimulation side. In the second test session, we repeated this procedure but assigned the other side of the chamber as the stimulation side. The behavior of the mice was recorded with a CCD camera interfaced with Ethovision software (v.11.5; Noldus Information Technologies), which was also used to control the laser stimulation and extract behavioral parameters (position, time, distance and velocity).

### Elevated plus maze test

The elevated plus maze (EPM) test consisted of a non-transparent, cross-shaped apparatus made of Plexiglass, with two ‘closed’ arms enclosed by 15-cm-high walls, and two ‘open’ arms without walls. The arms were 30-cm long and 5-cm wide and extended from a central platform (5 cm × 5 cm), allowing mice to freely move across the arms. The maze was elevated at a height of 55 cm from the ground. At the start of the 10-min sessions, mice were placed in the central platform. Animals’ behavior was videotaped, and the resulting data were analyzed using the image processing and tracking software Ethovision XT 5.1 (Noldus Information Technologies).

### Behavioral data acquisition and analysis

Behavior experiments were conducted with an open-source platform based on the Bpod State Machine (Sanworks). In the go/no-go task, licking data were acquired by a custom ‘lickometer’—a licking detection circuit composed of the metal spout, the mouse, and a ground wire connected to the tail of the mouse. Each time the mice licked the spout, the detection circuit was completed and a lick event was registered. The lick events were recorded by Bpod and saved on a computer. In the go/no-go task, the ‘hit’ rate was calculated as the number of hit trials divided by the total number of go trials and the ‘CR’ rate was calculated as the number of CR trials divided by the total number of no-go trials.

## QUANTIFICATION AND STATISTICAL ANALYSIS

All statistical tests are indicated where used. Statistical analyses were conducted using GraphPad Prism (v.7; GraphPad Software), MATLAB (2017a) statistical toolbox (MathWorks) and Igor Pro (WaveMetrics). To determine whether parametric tests could be used, the D’Agostino– Pearson test or Shapiro–Wilk test was performed on all data as a test for normality. Parametric tests were used whenever possible to test differences between two or more means. Non-parametric tests were used when data distributions were non-normal. Analysis of variance (ANOVA) was used to check for main effects and interactions in experiments with repeated measures and more than one factor. When main effects or interactions were significant, we performed the planned comparisons according to experimental design (for example, comparing laser on and off conditions). All comparisons were two-tailed. Statistical significance was set at the level of P < 0.05. All data are shown as mean ± standard error of the mean (SEM) unless stated otherwise.

## FIGURES and SUPPLEMENTARY FIGURES

**Supplementary Figure 1.**
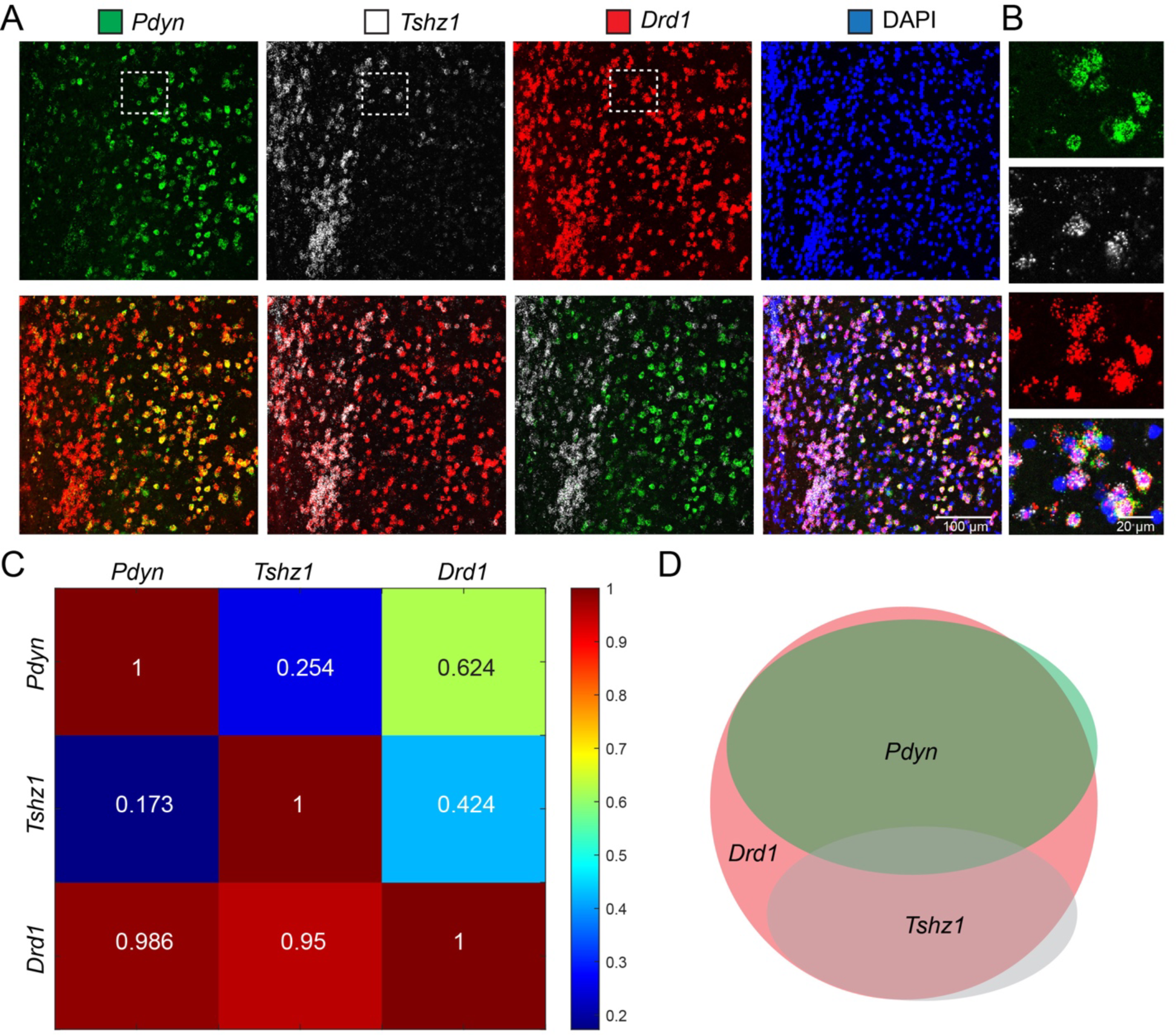
Characterization of the different subtypes of D1 neurons in the NAc. (A) Upper: confocal images of a coronal NAc section, showing the distribution of *Pdyn*, *Tshz1* and *Drd1* detected with smFISH. Lower, starting from the leftmost panel: overlay of the *Pdyn* and *Drd1*, overlay of the *Tshz1* and *Drd1*, overlay of the *Pdyn* and *Tshz1*, and overlay of all images in the upper panels. This experiment was repeated in 3 mice. (B) High-magnification images of the boxed area in A, showing that *Tshz1* and *Pdyn* do not overlap, but both overlap with *Drd1*. (C) Quantification of the overlap between neurons that express *Pdyn*, *Tshz1* and *Drd1* detected with smFISH in the NAc. Each value in parentheses represents the number of neurons detected with the respective probe in 3 mice. Values in matrix represent the percent labeling with probes in rows among neurons labeled with probes in columns. For example, 25.4% of *Tshz1*-labeled neurons were also labeled with *Pdyn*. (D) A schematic showing the relationship between different populations in the NAc.

**Supplementary Figure 2.**
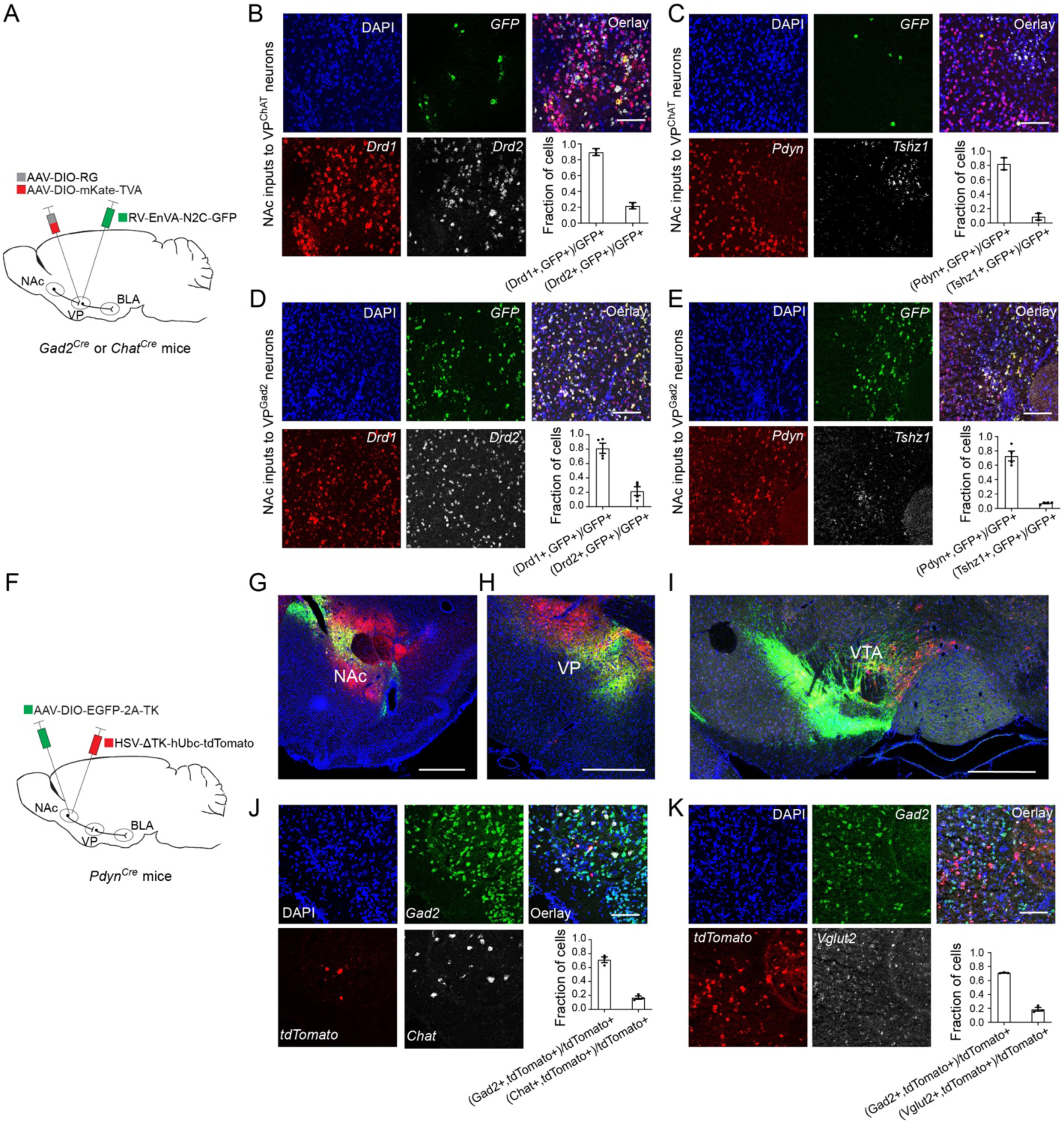
Mapping the connectivity between NAc neurons and VP neurons. (A) A schematic of the approach. (B and C) Identifying NAc neuronal types that provide monosynaptic inputs to VP^ChAT^ neurons. (B) Confocal images of a coronal NAc section, showing the distribution of *GFP*, *Drd1* and *Drd2* detected with smFISH. Lower-right panel: quantification of the fractions of rabies-labeled (*GFP*^+^) cells that express *Drd1* or *Drd2*. N = 2 mice. (C) Same as B, except that *GFP*, *Pdyn* and *Tshz1* were detected with smFISH. Lower-right panel: quantification of the fractions of rabies-labeled cells that express *Pdyn* or *Tshz1*. N = 2 mice. (D and E) Identifying NAc neuronal types that provide monosynaptic inputs to VP^Gad2^ neurons. (D) Confocal images of a coronal NAc section, showing the distribution of *GFP*, *Drd1* and *Drd2* detected with smFISH. Lower-right panel: quantification of the fractions of rabies-labeled (*GFP*^+^) cells that express *Drd1* or *Drd2*. N = 4 mice. (E) Same as D, except that *GFP*, *Pdyn* and *Tshz1* were detected with smFISH. Lower-right panel: quantification of the fractions of rabies-labeled cells that express *Pdyn* or *Tshz1*. N = 4 mice. (F) A schematic of the approach. (G-I) Confocal images showing the virus injection site (G) and the HSV-labeled (tdTomato^+^) neurons in the VP (H) and VTA (I), which were detected with immunohistochemistry. (J) Confocal images of a coronal VP section, showing the distribution of *Gad2*, *tdTomato* and *ChAT* detected with smFISH. Lower-right panel: quantification of the fractions of HSV-labeled cells that express *Gad2* or *ChAT*. N = 3 mice. (K) Same as J, except that *Gad2*, *tdTomato* and *Vglut2* were detected with smFISH. Lower-right panel: quantification of the fractions of HSV-labeled cells that express *Gad2* or *Vglut2*. N = 3 mice. Scale bars in B, C, D, E, J, and K: 100 µm. Scale bars in G-I: 1 mm. Data are presented as mean ± s.e.m.

**Supplementary Figure 3.**
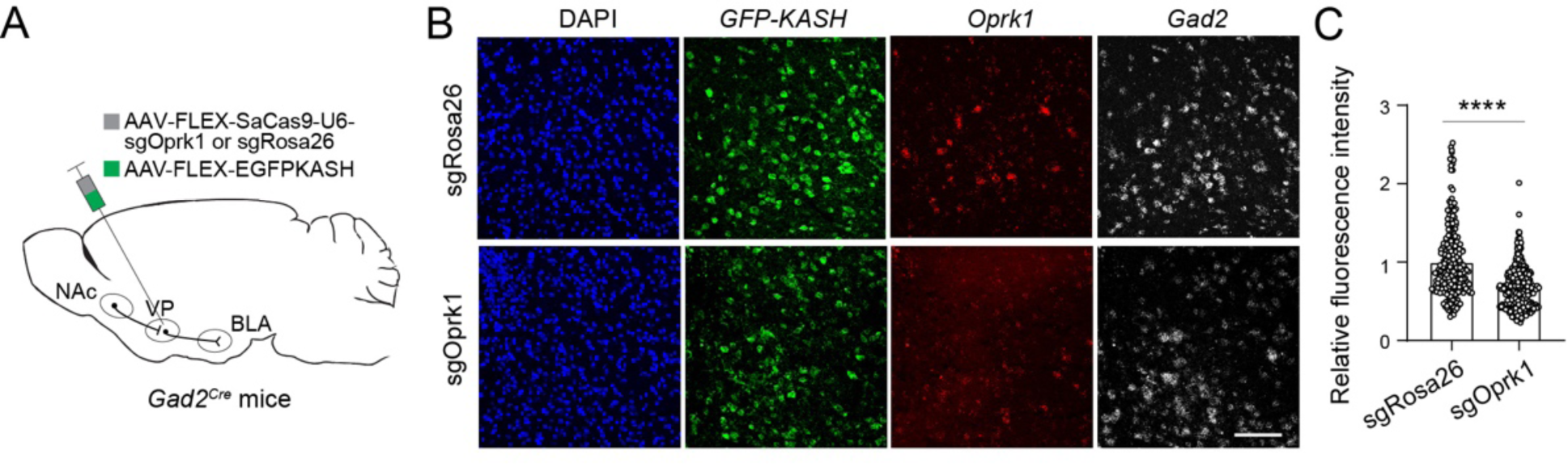
Suppressing *Oprk1* expression in VP GABAergic neurons with CRISPR/Cas9. (A) A schematic of the approach. (B) Confocal images of a coronal VP section, showing signals for *GFP-KASH*, *Oprk1* and *Gad2* detected with smFISH in a sgRosa26 control mouse (upper) and a sgOprk1 mouse (lower). Scale bar, 100 µm. (C) Quantification of relative fluorescent intensity of *Oprk1* mRNA signals in individual cells identified as *Oprk1*-positive. Data was normalized to the mean fluorescent intensity of the sgRosa26 group (sgRosa26, n = 278 cells from 3 mice; sgOprk1, n = 349 cells from 3 mice; t = 11.77, ****P < 1×10^-15^, unpaired t test). Data are presented as mean ± s.e.m.

**Supplementary Figure 4.**
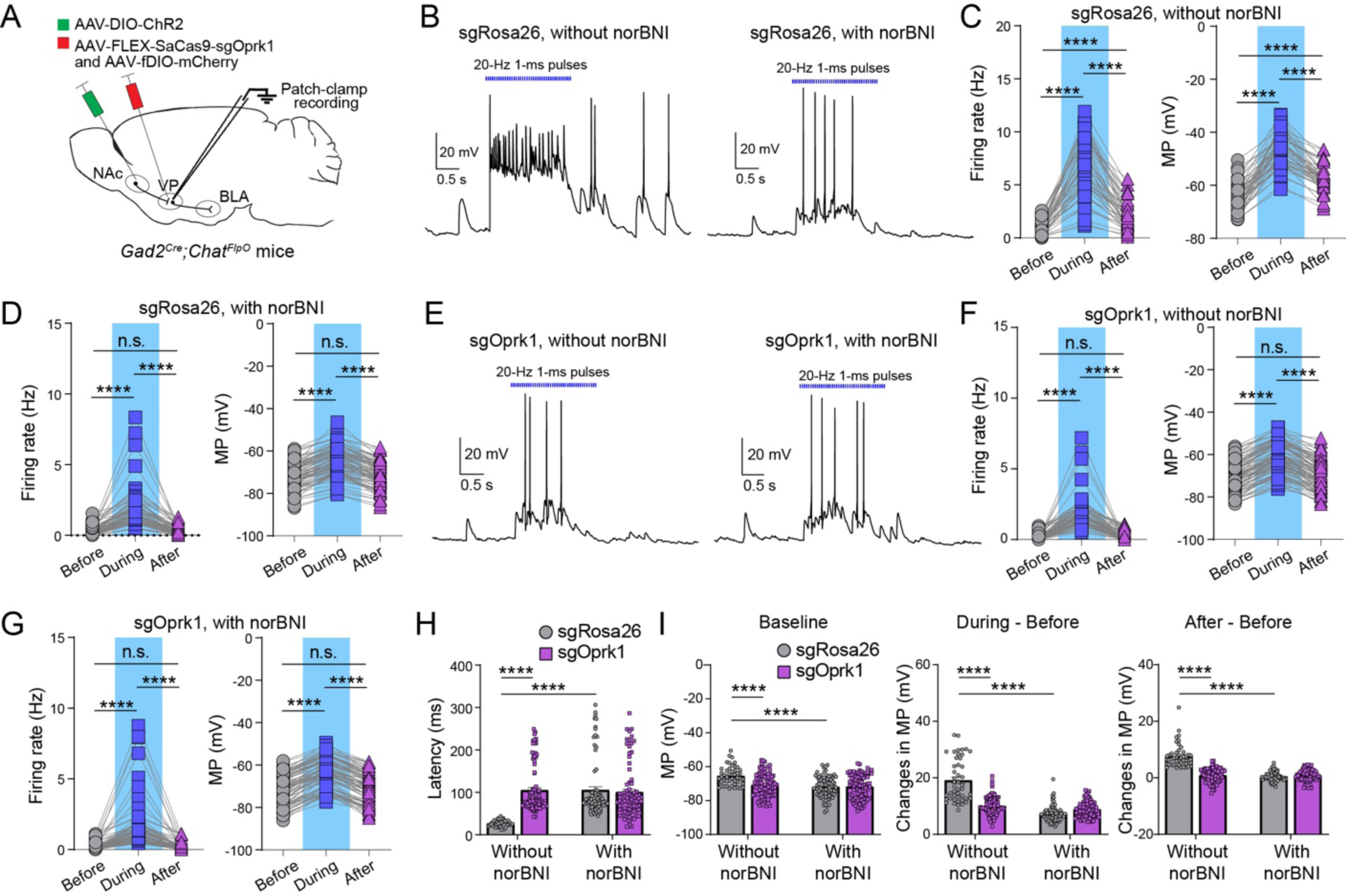
NAc neuron-driven disinhibition of VP^ChAT^ neurons depends on KORs in VP^Gad2^ neurons. (A) A schematic of the approach. (B) Traces of current-clamp recording from VP^ChAT^ neurons in response to photo-stimulation of axon terminals originating from NAc^Gad2^ neurons in sgRosa26 control mice. Left: without norBNI; right: with norBNI (100 nM). (C) Quantification of firing rate (left) and membrane potential (MP, right) before, during and after the photo-stimulation in the absence of norBNI (n = 46 neurons from 4 mice; firing rate, all ****P < 0.0001; MP, all ****P < 0.0001; Wilcoxon signed-rank test). (D) Same as C except that recording was performed in the presence of norBNI (n = 79 neurons from 4 mice; firing rate, ****P < 0.0001; n.s., nonsignificant, P = 0.5745; MP, ****P < 0.0001, n.s., P = 0.6023; Wilcoxon signed-rank test). (E) Traces of current-clamp recording from VP^ChAT^ neurons in response to photo-stimulation of axon terminals originating from NAc^Gad2^ neurons in sgOprk1 mice. Left: without norBNI; right: with norBNI (100 nM). (F) Quantification of firing rate (left) and MP (right) before, during and after the photo-stimulation in the absence of norBNI (n = 90 neurons from 4 mice; firing rate, ****P < 0.0001, n.s., P = 0.8587; MP, all ****P < 0.0001, n.s., P = 0.81; Wilcoxon signed-rank test). (G) Same as F except that recording was performed in the presence of norBNI (n = 82 neurons from 4 mice; firing rate, ****P < 0.0001; n.s., P = 0.8587; MP, ****P < 0.0001, n.s., P = 0.2696; Wilcoxon signed-rank test). (H) Quantification of the latency of light-evoked firing (without norBNI: sgRosa26, n = 46 neurons from 4 mice, sgOprk1, n = 90 neurons from 4 mice; with norBNI: sgRosa26, n = 79 neurons from 4 mice, sgOprk1, n = 82 neurons from 4 mice; F(1,126) = 53.34, ****P < 0.0001, two-way ANOVA followed by Sidak multiple comparisons test). (I) Quantification of baseline MP (left), changes in MP from baseline to photo-stimulation period (middle), and changes in MP from baseline to after photo-stimulation period (right) (without norBNI: sgRosa26, n = 46 neurons from 4 mice, sgOprk1, n = 90 neurons from 4 mice; with norBNI: sgRosa26, n = 79 neurons from 4 mice, sgOprk1, n = 82 neurons from 4 mice; left, F(1,126) = 22.93, ****P < 0.0001; middle, F(1,126) = 62.4, ****P < 0.0001; right, F(1,126) = 153.2, ****P < 0.0001; two-way ANOVA followed by Sidak multiple comparisons test). Data are presented as mean ± s.e.m.

**Supplementary Figure 5.**
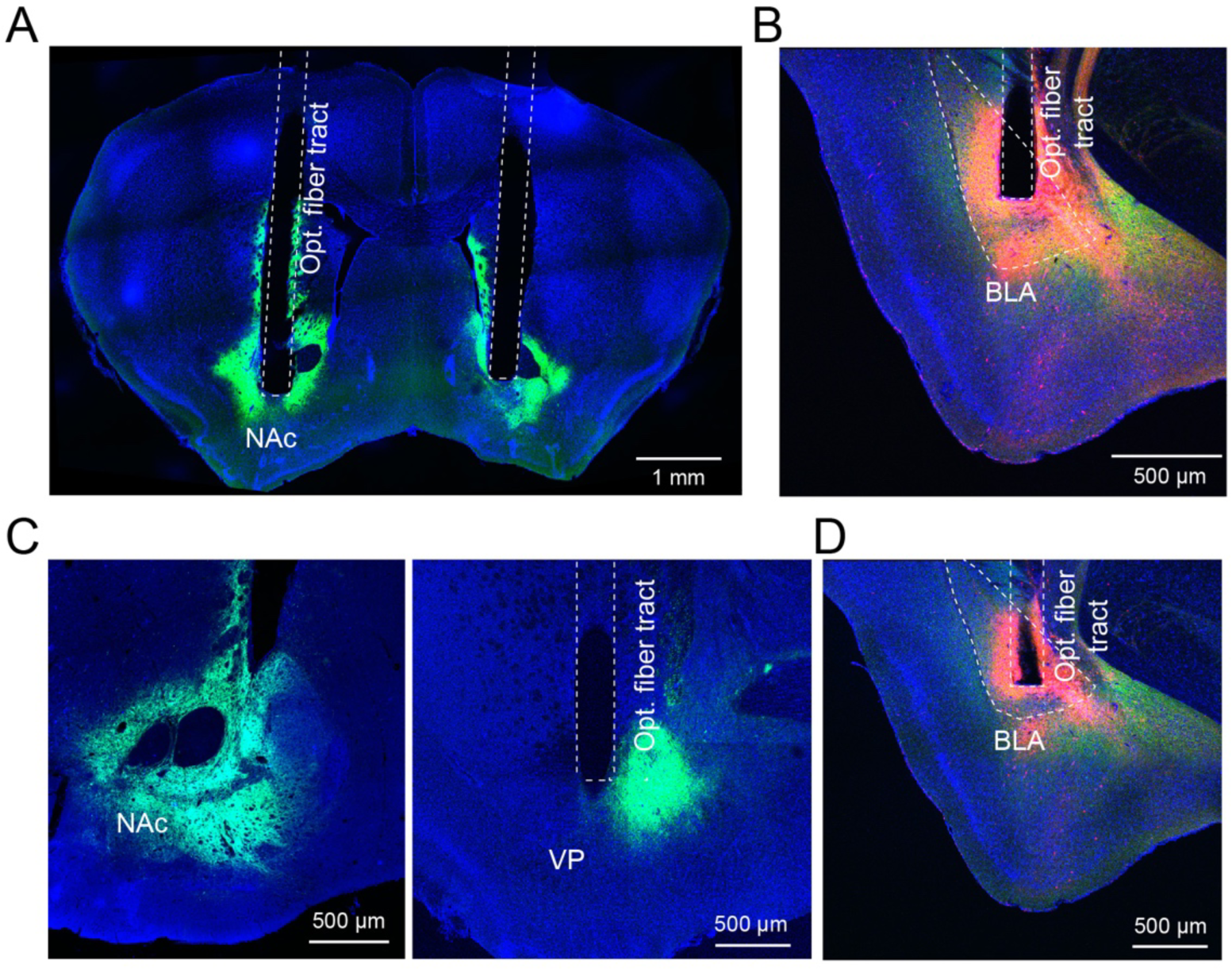
Histology for mice used in Figure 2 A-D. (A) A confocal image of a coronal brain section containing the NAc. The expression of ChR2-YFP in NAc^Pdyn^ neurons was detected by an antibody against green fluorescent protein (GFP). Optical fiber implantation locations in the NAc are indicated. (B) A confocal image of a coronal brain section containing the BLA in the same brain as in A. The expression of gACh3.0 (green) and mCherry (red) in the BLA was detected by antibodies against GFP and RFP (red fluorescent protein), respectively. The optical fiber implantation location in the BLA is indicated. (C) Confocal images of coronal brain sections containing the NAc (left) and VP (right). The expression of ChR2-YFP in NAc^Pdyn^ neurons (left) and their axon fibers in the VP (right) was recognized by an antibody against GFP. The optical fiber implantation location in the VP (right) is indicated. (D) A confocal image of a coronal brain section containing the BLA in the same brain as in C. The expression of gACh3.0 (green) and mCherry (red) in the BLA is detected by antibodies against GFP and RFP, respectively. The optical fiber implantation location in the BLA is indicated.

**Supplementary Figure 6.**
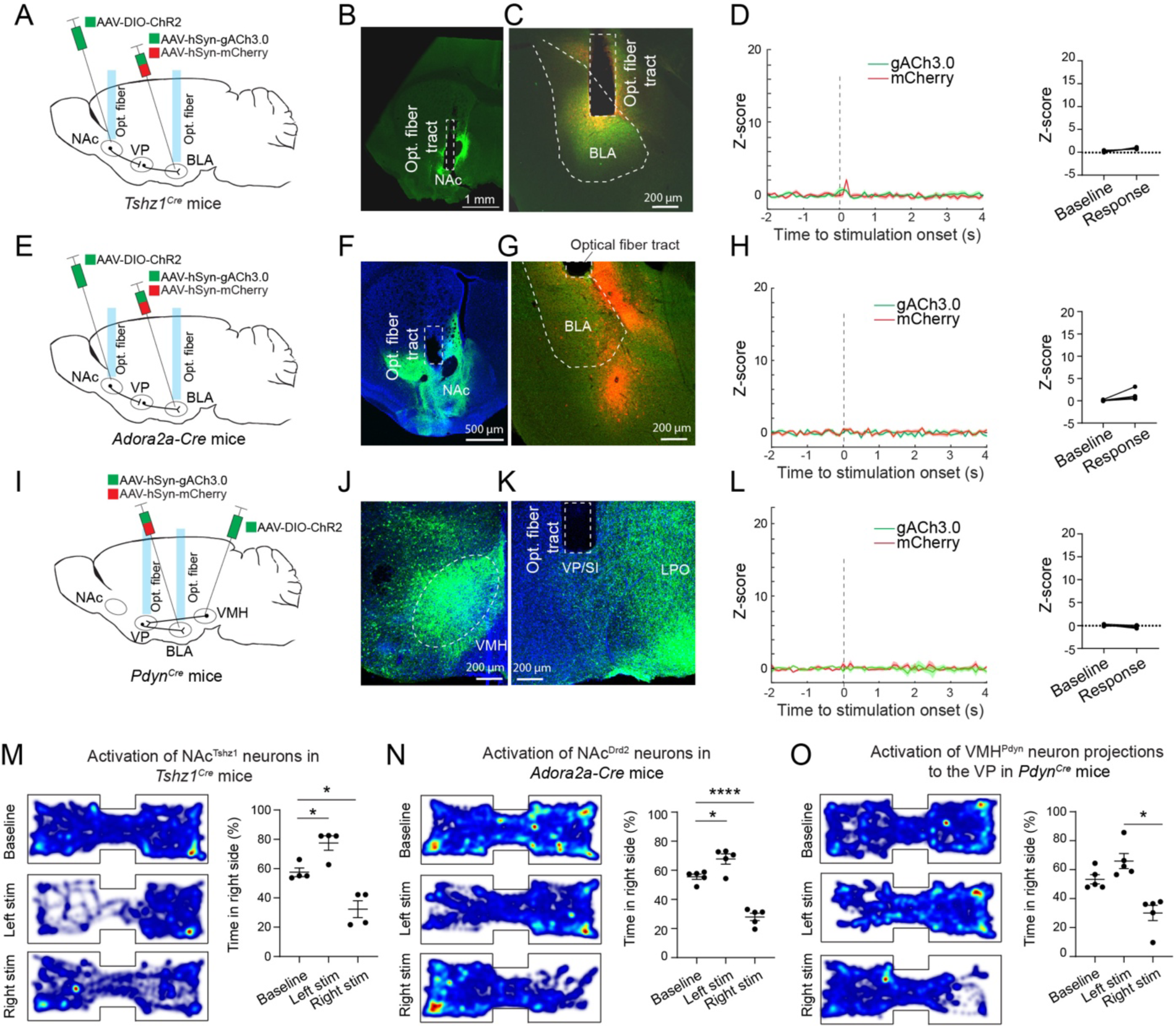
The effects of activating different neuronal types or pathways. (A) A schematic of the approach. (B) A confocal image of a coronal brain section containing the NAc. The expression of ChR2-YFP in NAc^Tshz1^ neurons was detected by an antibody against GFP. The optical fiber implantation location in the NAc is indicated. (C) A confocal image of a coronal brain section containing the BLA in the same brain as in B. The expression of gACh3.0 (green) and mCherry (red) in the BLA was detected by antibodies against GFP and RFP, respectively. The optical fiber implantation location in the BLA is indicated. (D) Left: average gACh3.0 signals from a mouse that received photo-stimulation of NAc^Tshz1^ neurons. mCherry signals are also shown to monitor potential motion artifacts. Right: quantification of the photo- stimulation-evoked response for all mice. Only minor response can be detected (n = 4 mice, t = 3.24, P = 0.0478, paired t test). (E) A schematic of the approach. (F) A confocal image of a coronal brain section containing the NAc. The expression of ChR2-YFP in NAc^Drd2^ neurons was detected by an antibody against GFP. The optical fiber implantation location in the NAc is indicated. (G) A confocal image of a coronal brain section containing the BLA in the same brain as in F. The expression of gACh3.0 (green) and mCherry (red) in the BLA was detected by antibodies against GFP and RFP, respectively. The optical fiber implantation location in the BLA is indicated. (H) Left: average gACh3.0 signals from a mouse that received photo-stimulation of NAc^Drd2^ neurons. mCherry signals are also shown to monitor potential motion artifacts. Right: quantification of the photo- stimulation-evoked response for all mice. No response can be detected (n = 5 mice, t = 2.51, P = 0.066, paired t test). (I) A schematic of the approach. (J, K) Confocal images of coronal brain sections containing the VMH (J) and the VP (K). The expression of ChR2-YFP in VMH^Pdyn^ neurons (J) and their axons terminals in the VP (K) was detected by an antibody against GFP. The optical fiber implantation location in the VP is indicated. (L) Left: average gACh3.0 signals in the BLA from a mouse that received photo-stimulation of VMH^Pdyn^ neuron terminals in the VP. mCherry signals are also shown to monitor potential motion artifacts. Right: quantification of the photo-stimulation-evoked response for all mice. No response can be detected (n = 5 mice, t = 2.04, P = 0.11, paired t test). (M) Left: heatmaps of the activity of a mouse at baseline (top) and receiving photo-stimulation of NAc^Tshz1^ neurons once entering the left (middle) or right (bottom) side of the chamber. Right: quantification of the time spent in the right side of the chamber under different situations (n = 4 mice, F = 23.47, P = 0.0003, *P = 0.035, *P = 0.0104, one-way ANOVA followed by Tukey’s multiple comparisons test). (N) Left: heatmaps of the activity of a mouse at baseline (top) and receiving photo-stimulation of NAc^Drd2^ neurons once entering the left (middle) or right (bottom) side of the chamber. Right: quantification of the time spent in the right side of the chamber under different conditions (n = 5 mice, F = 57.25, P = 7.3×10^-7^, *P = 0.019, ****P = 6.25×10^-7^, one-way ANOVA followed by Tukey’s multiple comparisons test). (O) Left: heatmaps of the activity of a mouse at baseline (top) and receiving photo-stimulation of VMH^Pdyn^ neuron projections to the VP once entering the left (middle) or right (bottom) side of the chamber. Right: quantification of the time spent in the right side of the chamber under different conditions (n = 5 mice, F = 15.3, P = 0.0005, *P = 0.0105, one-way ANOVA followed by Tukey’s multiple comparisons test). Data are presented as mean ± s.e.m.

**Supplementary Figure 7.**
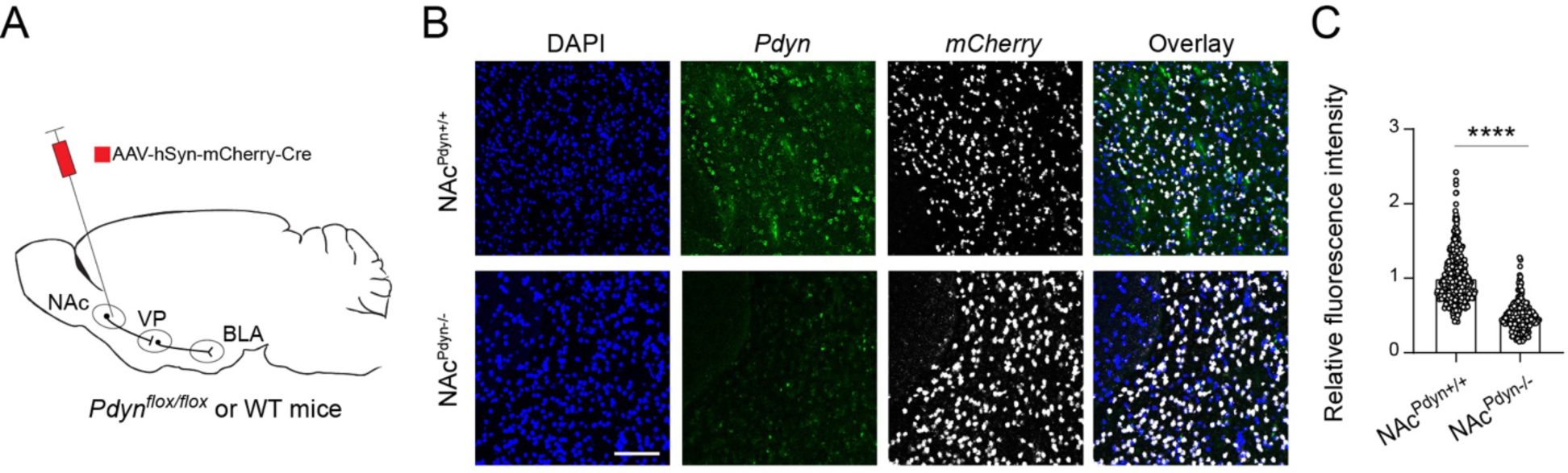
Deletion of *Pdyn* in NAc neurons. (A) A schematic of the approach. (B) Confocal images of a coronal NAc section, showing signals for *Pdyn* and *mCherry* detected with smFISH in a NAc^Pdyn+/+^ control mouse (upper) and a NAc^Pdyn-/-^ mouse (lower). Scale bar, 100 µm. (C) Quantification of relative fluorescent intensity of *Pdyn* mRNA signals in individual cells identified as *mCherry*-positive. Data was normalized to the mean fluorescent intensity of control group (NAc^Pdyn+/+^, n = 495 cells from 3 mice; NAc^Pdyn-/-^, n = 286 cells from 3 mice; t = 24.34, ****P < 1×10^-15^, unpaired t test). Data are presented as mean ± s.e.m.

**Supplementary Figure 8.**
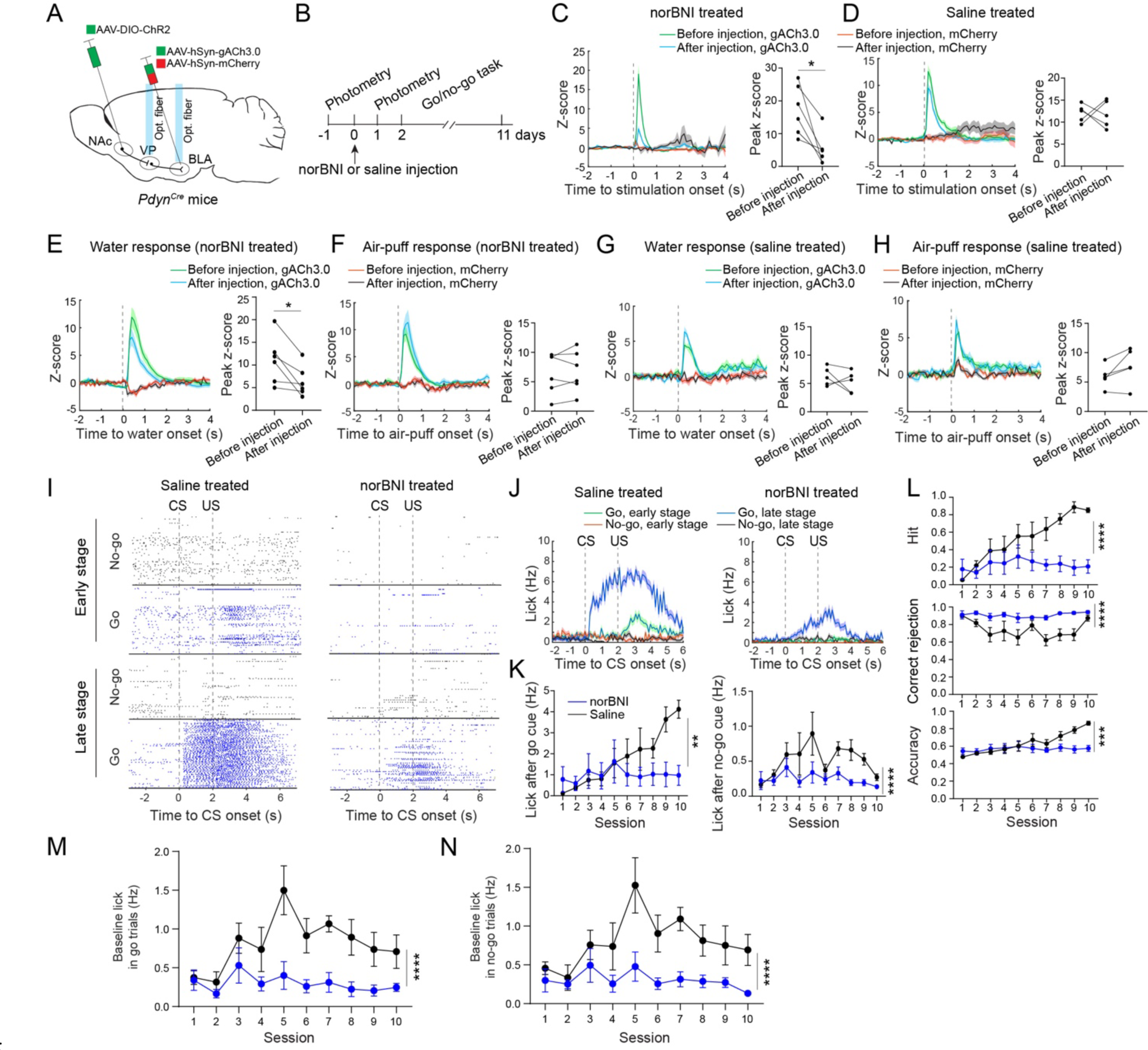
Dynorphin/KOR signaling is required for acetylcholine release in the BLA and reward-seeking behavior in mice. (A, B) A schematic of the approach (A) and experimental design (B). (C) Left: average gACh3.0 signals from a mouse that received photo-stimulation of NAc^Pdyn^®VP projections, before and after systemic administration of norBNI. mCherry signals are also shown to monitor potential motion artifacts. Right: quantification of the photo-stimulation-evoked response for all mice (n = 6 mice, t = 3.85, *P = 0.012, paired t test). (D) Same as C, except that the data were from mice received systemic administration of saline instead of norBNI (n = 5 mice, t = 0.086, P = 0.935, paired t test). (E) Left: average gACh3.0 signals from a mouse that received water, before and after systemic administration of norBNI. mCherry signals are also shown to monitor potential motion artifacts. Right: quantification of the response to water for all mice (n = 6 mice, t = 3.3, *P = 0.0218, paired t test). (F) Left: average gACh3.0 signals from a mouse that received air-puff, before and after systemic administration of norBNI. mCherry signals are also shown to monitor potential motion artifacts. Right: quantification of the response to air-puff for all mice (n = 6 mice, t = 0.64, P = 0.55, paired t test). (G) Same as E, except that the data were from mice received systemic administration of saline instead of norBNI (n = 5 mice, t = 0.924, P = 0.4, paired t test). (H) Same as F, except that the data were from mice received systemic administration of saline instead of norBNI (n = 5 mice, t = 2.23, P = 0.09, paired t test). (I) Lick raster of a saline-treated mouse (left) and a nor-BNI-treated mouse (right) during go/no-go training. (J) Average licking rates of the same mice in I. (K) Quantification of licking rates following CS onset in go trials (left) and no-go trials (right) across training sessions (go trials: F(1,100) = 8.4, **P = 0.0046; no-go trials: F(1,100) = 18.92, ****P = 0.00003; norBNI group, n = 6 mice, saline group, n = 6 mice, two-way ANOVA followed by Sidak’s test). (L) Top: hit rate, F(1,100) = 41.52, ****P = 4.13×10^-9^. Middle: CR rate, F(1,100) = 48.42, ****P = 3.64×10^-10^. Bottom: accuracy, F(1,100) = 13.3, ***P = 0.0004. norBNI group, n = 6 mice, saline group, n = 6 mice, two-way ANOVA followed by Sidak’s test. (M and N) Baseline licking rates during go/no-go training (M: go trials, F(1,100) = 43.92, ****P = 1.75×10^-9^; N: no-go trials, F(1,100) = 38.46, ****P = 1.27×10^-8^; norBNI group, n = 6 mice, saline group, n = 6 mice, two-way ANOVA followed by Sidak’s test). Data are presented as mean ± s.e.m.

**Supplementary Figure 9.**
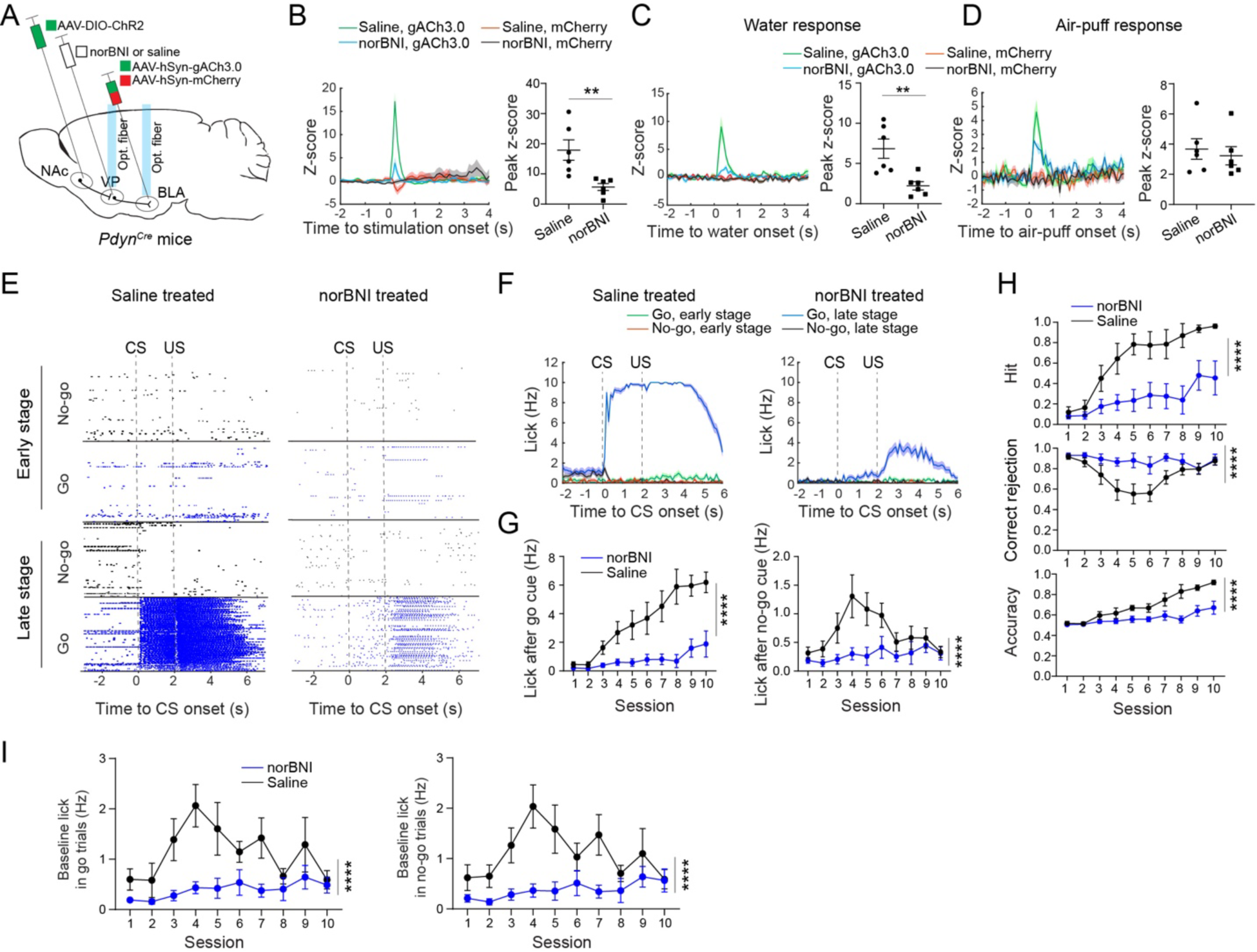
Dynorphin/KOR signaling in the VP is required for acetylcholine release in the BLA and reward-seeking behavior in mice. (A) A schematic of the approach. (B) Left: average gACh3.0 signals from each of two mice that received local injection of either norBNI or saline into the VP. Both mice received photo-stimulation of NAc^Pdyn^®VP projections. mCherry signals are also shown to monitor potential motion artifacts. Right: quantification of the photo-stimulation-evoked response for all mice (n = 6 mice in each group, t = 3.44, **P = 0.0063, unpaired t test). (C) Left: average gACh3.0 signals from each of two mice that received local injection of either norBNI or saline into the VP. Both mice received water. mCherry signals are also shown. Right: quantification of the response to water for all mice (n = 6 mice for each group, t = 3.52, **P = 0.0056, unpaired t test). (D) Same as C, except that the data were from mice received air-puff instead of water (quantification of the response to air-puff, n = 6 mice in each group, t = 0.5, P = 0.63, unpaired t test). (E) Lick raster of a saline-treated mouse (left) and a nor-BNI-treated mouse (right) during go/no-go training. (F) Average licking rates of the same mice in E. (G) Quantification of licking rates following CS onset in go trials (left) and no-go trials (right) across training sessions (go trials: F(1,110) = 90.95, ****P < 1×10^-15^; no-go trials: F(1,110) = 26.77, ****P = 1×10^-6^; norBNI group, n = 7 mice, saline group, n = 6 mice, two-way ANOVA followed by Sidak’s test. (H) Top: hit rate, F(1,110) = 66.07, ****P = 7.1×10^-13^. Middle: CR rate, F(1,110) = 26.24, ****P = 1.29×10^-6^. Bottom: accuracy, F(1,110) = 56.91, ****P = 1.4×10^-11^. norBNI group, n = 7 mice, saline group, n = 6 mice, two-way ANOVA followed by Sidak’s test. (I) Baseline licking rates during go/no-go training (left: go trials, F(1,110) = 37.76, ****P = 1.31×10^-8^; right: no-go trials, F(1,110) = 38.69, ****P = 9.2×10^-9^; norBNI group, n = 7 mice, saline group, n = 6 mice, two-way ANOVA followed by Sidak’s test). Data are presented as mean ± s.e.m.

**Supplementary Figure 10.**
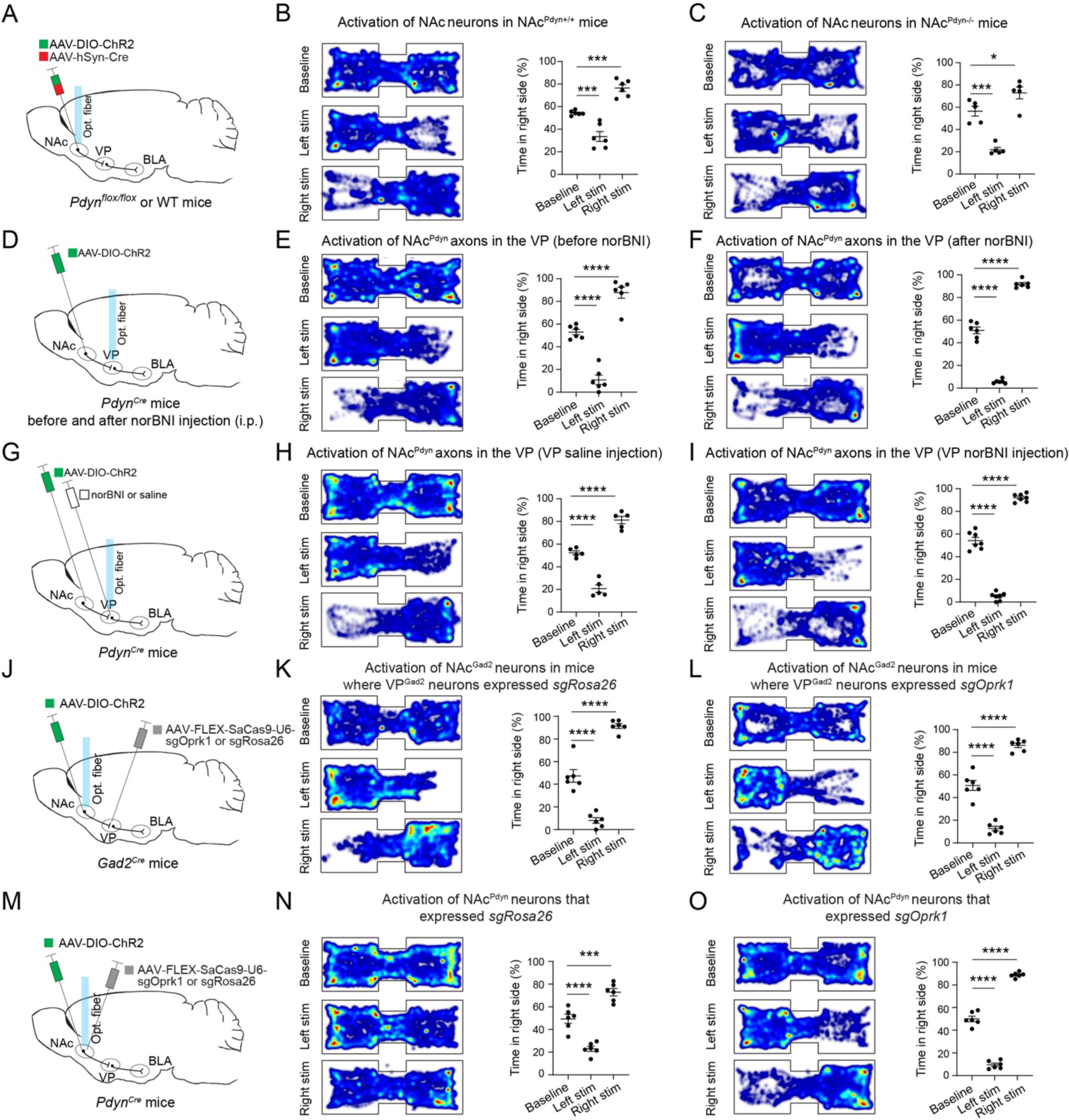
Manipulation of dynorphin/KOR signaling does not affect place preference induced by photo-stimulating the NAc®VP pathway. (A) A schematic of the approach. (B) Data from mice that had intact *Pdyn* in NAc neurons (NAc^Pdyn+/+^ mice). Left: heatmaps of the activity of a mouse at baseline (top) and receiving photo-stimulation of NAc neurons once entering the left (middle) or right (bottom) side of the chamber. Right: quantification of the time spent in the right side of the chamber under different situations (n = 6 mice, F = 48.28, P = 2.9×10^-7^, ***P = 0.0007, ***P = 0.0004, one-way ANOVA followed by Tukey’s multiple comparisons test). (C) Same as B, except that data were from mice where *Pdyn* in NAc neurons was deleted (NAc^Pdyn-/-^ mice) (n = 5 mice, F = 38.6, P = 5.93×10^-6^, ***P = 0.0002, *P = 0.042, one-way ANOVA followed by Tukey’s multiple comparisons test). (D) A schematic of the approach. (E) Data from mice before receiving systemic administration of norBNI. Left: heatmaps of the activity of a mouse at baseline (top) and receiving photo-stimulation of NAc^Pdyn^ neuron projections to the VP once entering the left (middle) or right (bottom) side of the chamber. Right: quantification of the time spent in the right side of the chamber under different situations (n = 6 mice, F = 95.1, P = 3×10^-9^, ****P = 5×10^-6^, ****P = 4.58×10^-5^, one-way ANOVA followed by Tukey’s multiple comparisons test). (F) Same as E, except that data were from mice after receiving systemic administration of norBNI (n = 6 mice, F = 503.3, P = 1.8×10^-14^, ****P = 1.2×10^-10^, ****P = 5.2×10^-10^, one-way ANOVA followed by Tukey’s multiple comparisons test). (G) A schematic of the approach. (H) Data from mice that received local injection of saline in the VP. Left: heatmaps of the activity of a mouse at baseline (top) and receiving photo-stimulation of NAc^Pdyn^ neuron projections to the VP once entering the left (middle) or right (bottom) side of the chamber. Right: quantification of the time spent in the right side of the chamber under different situations (n = 5 mice, F = 112.1, P = 1.7×10^-8^, ****P = 0.000013, ****P = 0.00003, one-way ANOVA followed by Tukey’s multiple comparisons test). (I) Same as H, except that data were from mice that received local injection of norBNI in the VP (n = 6 mice, F = 470.6, P < 1×10^-15^, ****P = 3.3×10^-12^, ****P = 2.7×10^-10^, one-way ANOVA followed by Tukey’s multiple comparisons test). (J) A schematic of the approach. (K) Data from mice in which VP^Gad2^ neurons expressed the control guide RNA *sgRosa26*. Left: heatmaps of the activity of a mouse at baseline (top) and receiving photo-stimulation of NAc^Gad2^ neurons once entering the left (middle) or right (bottom) side of the chamber. Right: quantification of the time spent in the right side of the chamber under different situations (n = 6 mice, F = 124.6, P = 4.5×10^-10^, ****P = 5.78×10^-6^, ****P = 1.49×10^-6^, one-way ANOVA followed by Tukey’s multiple comparisons test). (L) Same as K, except that data were from mice in which VP^Gad2^ neurons expressed the guide RNA *sgOprk1* (n = 6 mice, F = 146.2, P = 1.45×10^-10^, ****P = 7.05×10^-7^, ****P = 1.65×10^-6^, one-way ANOVA followed by Tukey’s multiple comparisons test). (M) A schematic of the approach. (N) Data from mice in which NAc^Pdyn^ neurons expressed the control guide RNA *sgRosa26*. Left: heatmaps of the activity of a mouse at baseline (top) and receiving photo-stimulation of NAc^Pdyn^ neurons once entering the left (middle) or right (bottom) side of the chamber. Right: quantification of the time spent in the right side of the chamber under different situations (n = 6 mice, F = 60.56, P = 6.5×10^-8^, ****P = 0.0000967, ***P = 0.0003, one-way ANOVA followed by Tukey’s multiple comparisons test). (O) Same as N, except that data were from mice in which NAc^Pdyn^ neurons expressed the guide RNA *sgOprk1* (n = 6 mice, F = 519.9, P = 1.4×10^-14^, ****P = 1.42×10^-10^, ****P = 2.64×10^-10^, one-way ANOVA followed by Tukey’s multiple comparisons test). Data are presented as mean ± s.e.m.

**Supplementary Figure 11.**
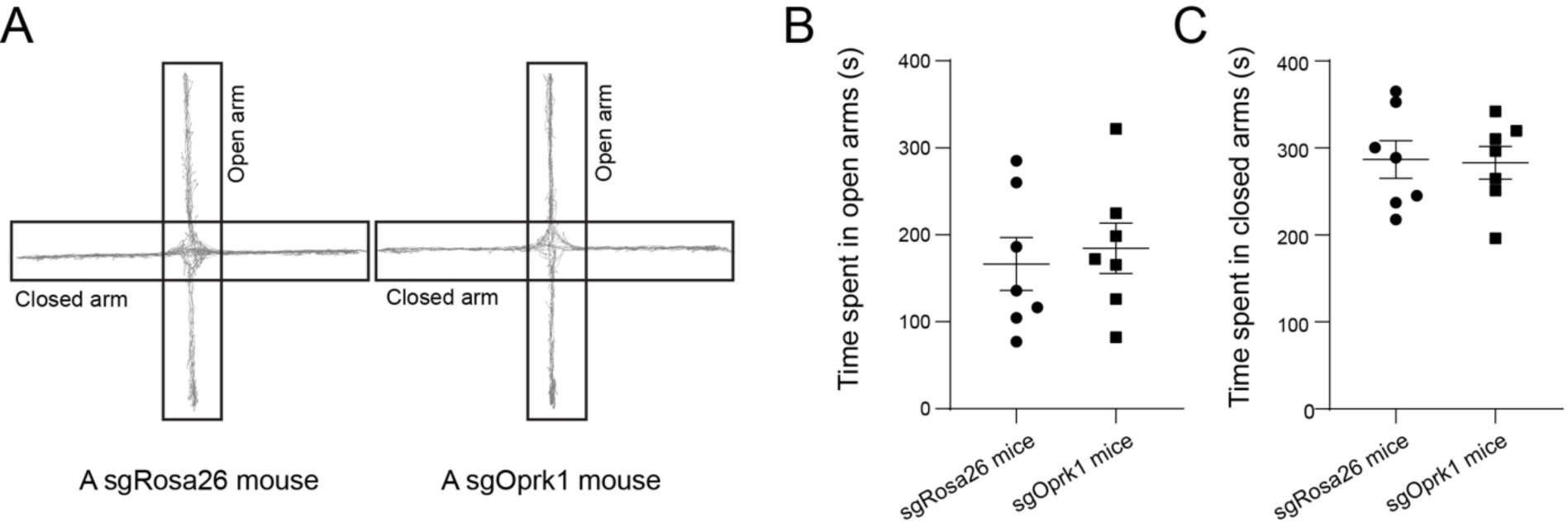
Suppressing KOR expression in VP GABAergic neurons does not affect anxiety levels. (A) Activity traces of a sgRosa26 mouse (left) and a sgOprk1 mouse in an elevated plus maze. (B, C) Quantification of time spent in the open arms (B) and closed arms (C) (n = 7 mice in each group; open arms, t = 0.43, P = 0.675; closed arms, t = 0.13, P = 0.9; unpaired t test). Data are presented as mean ± s.e.m.

**Supplementary Figure 12.**
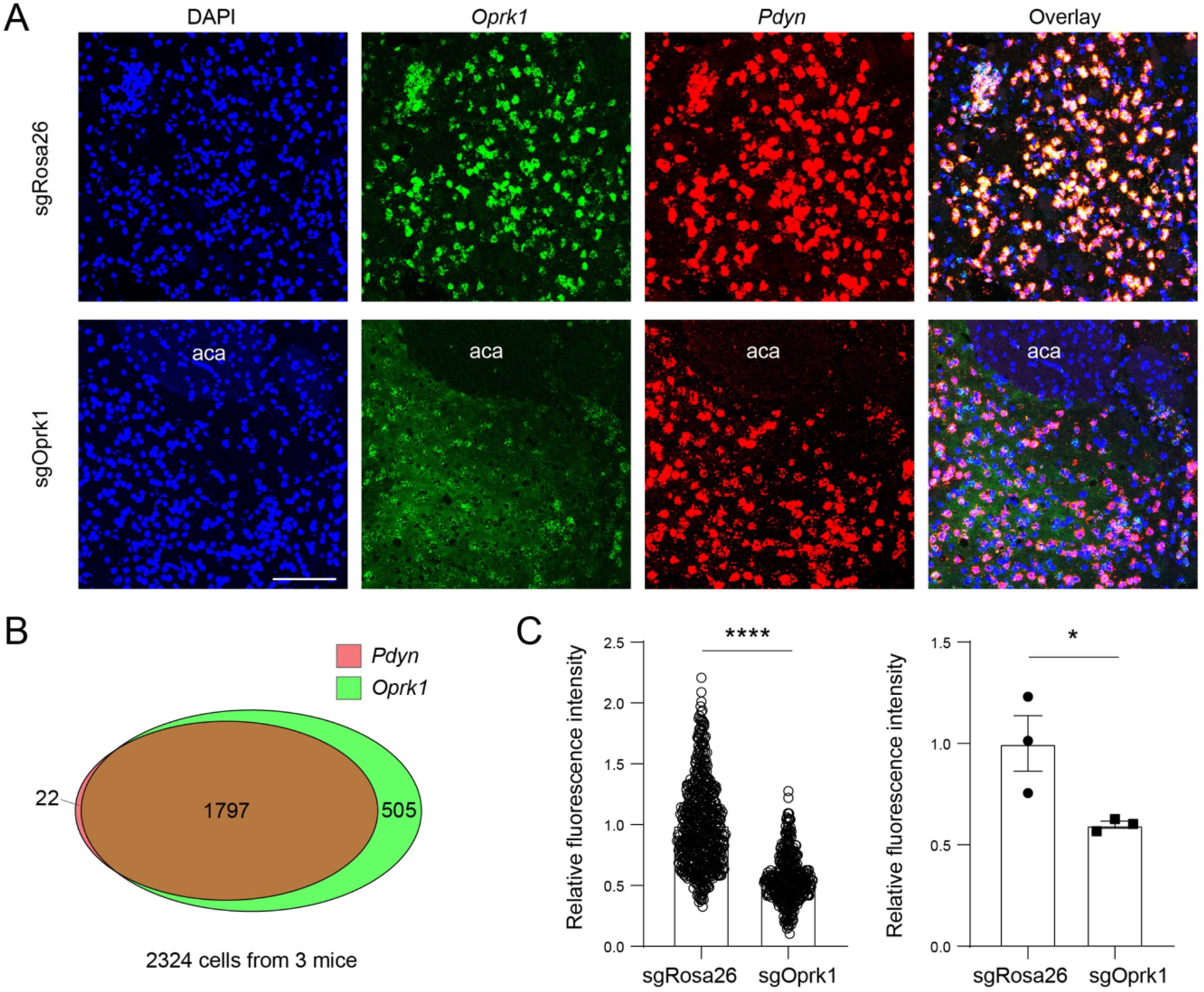
Suppressing *Oprk1* expression in NAc^Pdyn^ neurons with CRISPR/Cas9. (A) Confocal images of coronal NAc sections, showing signals for *Oprk1* and *Pdyn* detected with smFISH in a sgRosa26 control mouse (upper) and a sgOprk1 mouse (lower). Scale bar, 100 µm. (B) A schematic showing the relationship between *Pdyn*-expressing cells and *Oprk1*-expressing cells in the NAc. (C) Left: quantification of relative fluorescent intensity of *Oprk1* mRNA signals in individual cells identified as *Pdyn*-positive in the NAc. Data was normalized to the mean fluorescent intensity of sgRosa26 group (sgRosa26, n = 587 cells from 3 mice; sgOprk1, n = 385 cells from 3 mice; t = 22.56, ****P <1×10^-15^, unpaired t test). Right: the statistics for individual mice (t = 2.9, *P = 0.0441, unpaired t test). Data are presented as mean ± s.e.m.

**Supplementary Figure 13.**
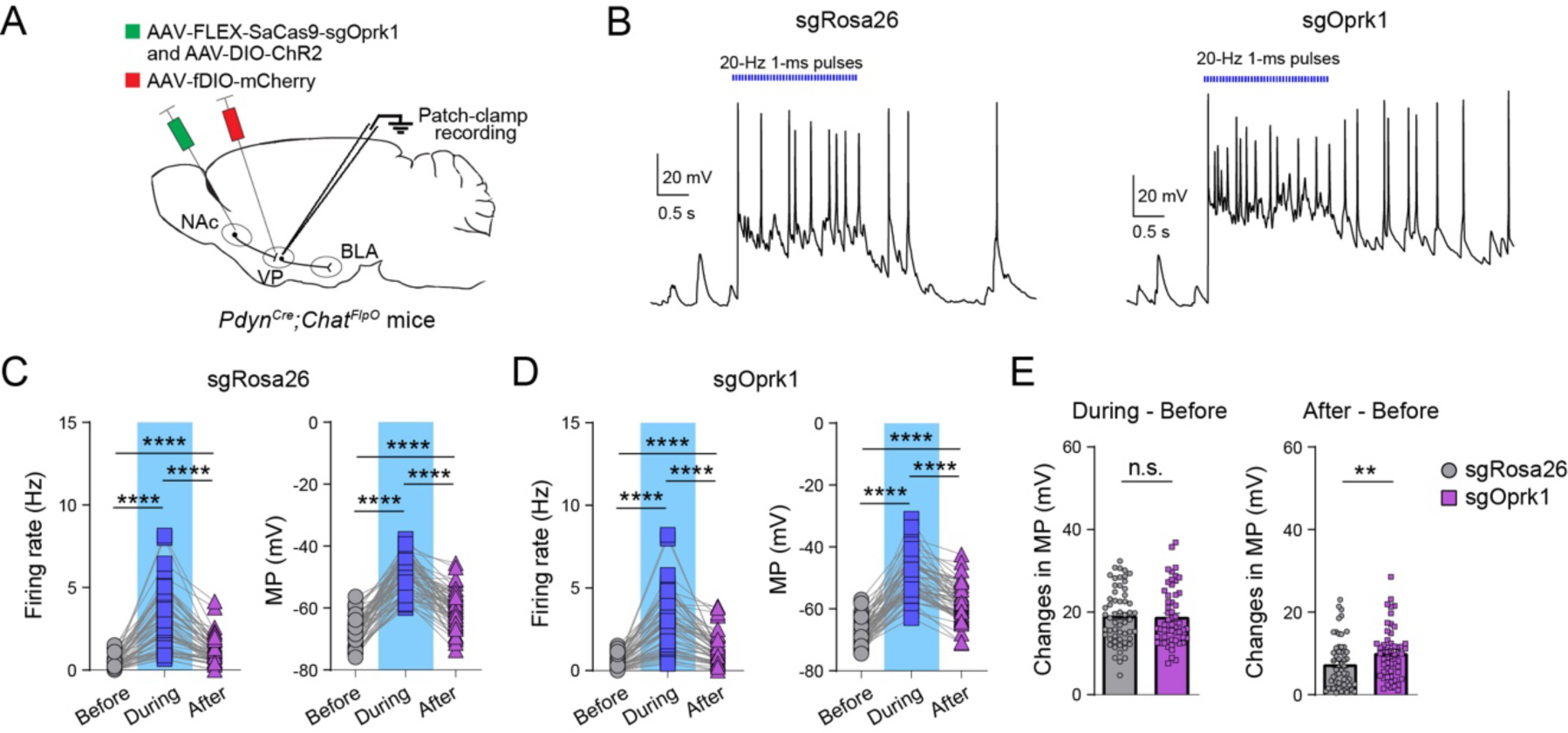
KORs in NAc^Pdyn^ neurons limit the disinhibition of VP^ChAT^ neurons. (A) A schematic of the approach. (B) Traces of current-clamp recording from cholinergic neurons in response to photo-stimulation of axon terminals originating from NAc^Pdyn^ neurons. Data are from a sgRosa26 control mouse (left) and a sgOprk1 mouse (right). (C) Quantification of firing rate (left) and membrane potential (MP, right) before, during and after the photo-stimulation in sgRosa26 control mice (n = 57 neurons from 3 mice; firing rate, ****P < 0.0001; MP, ****P < 0.0001, Wilcoxon signed-rank test). (D) Same as C except that recording was performed in sgOprk1 mice (n = 55 neurons from 3 mice; firing rate, ****P < 0.0001; MP, ****P < 0.0001, Wilcoxon signed-rank test). (E) Quantification of changes in MP from baseline to photo-stimulation period (left) and changes in MP from baseline to after photo-stimulation period (right) (sgRosa26, n = 57 neurons from 3 mice; sgOprk1, n = 55 neurons from 3 mice; n.s., nonsignificant, P = 0.7312; **P = 0.007, Mann-Whitney test). Data are presented as mean ± s.e.m.

**Supplementary Figure 14.**
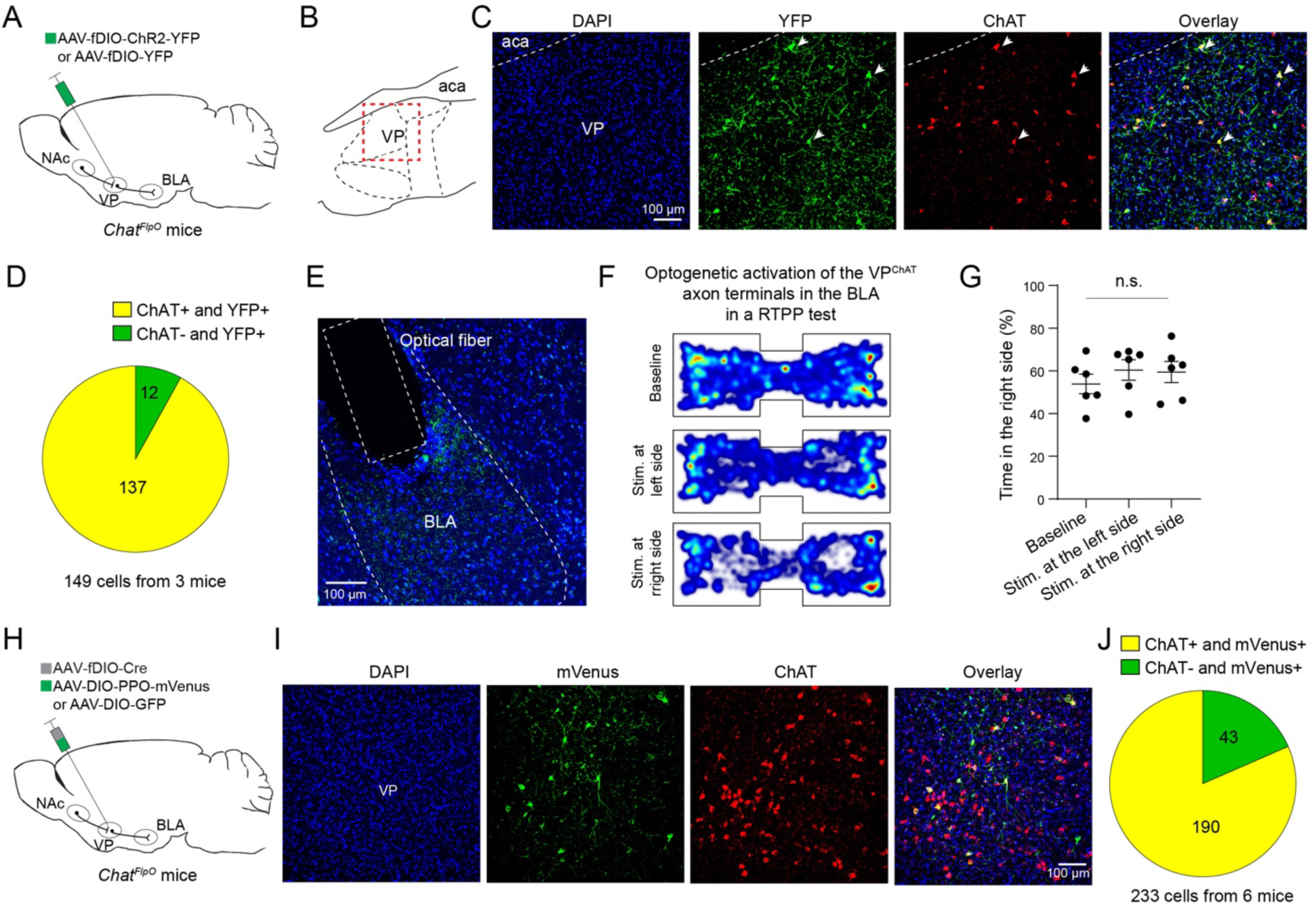
Targeting VP^ChAT^ neurons. (A) A schematic of the approach. (B) A diagram showing the position of VP area. The red box indicated the area imaged in C. (C) Confocal images of the virus injection site in the VP, showing the AAV-labeled (GFP^+^) neurons and ChAT-expressing (ChAT^+^) neurons, which were detected with immunohistochemistry. (D) Quantification of the overlap between the GFP^+^ neurons and the ChAT^+^ neurons. (E) A confocal image of a coronal brain section containing the BLA. The ChR2^+^ fibers (green) originating from VP^ChAT^ neurons were detected with an antibody against GFP. The optical fiber implantation location in the BLA is indicated. (F) Heatmaps of the activity of a mouse at baseline (top) and receiving photo-stimulation of VP^ChAT^ neuron projections to the BLA once entering the left (middle) or right (bottom) side of the chamber. (G) Quantification of the time spent in the right side of the chamber under different situations (n = 6 mice, F = 0.55, P = 0.588, one-way ANOVA followed by Tukey’s multiple comparisons test). (H) A schematic of the approach. (I) Confocal images of the virus injection site in the VP, showing the AAV-labeled (mVenus^+^) neurons and ChAT^+^ neurons, which were detected with immunohistochemistry. (J) Quantification of the overlap between the mVenus^+^ neurons and the ChAT^+^ neurons. Data are presented as mean ± s.e.m.

